# Dissolved inorganic carbon supports robust anabolism and methanogenesis in actively serpentinizing rocks

**DOI:** 10.1101/2025.08.25.672207

**Authors:** Tristan A. Caro, Srishti Kashyap, Ashley E. Maloney, David W. Hoyt, Michael D. Y. Kubo, Tori M. Hoehler, Alexis S. Templeton

**Author notes:** **Corresponding Authors:** Tristan A. Caro, Alexis S. Templeton. **Competing Interest Statement:** AST is currently a consulting geochemist at Eden Geopower Inc, which engages in geological hydrogen technology development.

## Abstract

Serpentinites, hydrated ultramafic rocks that produce [hyper]alkaline, reducing, H_2_-rich groundwaters, host subsurface microbial ecosystems. Though in the presence of enormous reducing power, life in serpentinizing systems is limited by oxidant and carbon availability. The forms of carbon that support the serpentinite-hosted microbiome, and their rates of biological assimilation, remain poorly understood. In this work, we quantify the habitability of subsurface environments shaped by serpentinization and examine the forms of carbon that support their microbial constituents, focusing specifically on dissolved inorganic carbon, acetate, and formate. We access reacted groundwater from Earth’s largest terrestrial serpentinizing body and measure carbon assimilation at the single-cell level. Across all conditions, we consistently observe robust assimilation of dissolved inorganic carbon into microbial biomass. Notably, we find that dissolved inorganic carbon supports the majority of methanogenic activity in the system, even at hyperalkaline conditions (pH > 11). Inferred bioenergetic fluxes suggest that rates of biological hydrogen-consumption and methanogenesis are relevant at the landscape scale. We identify a strong potential for the microbiome to be stimulated by increases in H_2_ and CO_2_, a finding with implications for the search for life on other planetary bodies and for the growing deployment of fluid injection technologies in ultramafic rocks, such as geological hydrogen production or carbon mineralization.

**Significance Statement:** When iron-rich rocks interact with water, they undergo a “serpentinization” reaction that results in the production of H_2_ and [hyper]alkaline fluids. This abiotic process does not operate alone but rather supports and interacts with microbial life. However, it remains unknown to what extent the stressors of subsurface life under [hyper]alkaline conditions may limit microbial activity. In this work, we quantify the rates at which microorganisms transform different carbon sources across a range of serpentinizing conditions. We find that serpentinizing rocks host a remarkably robust microbiome, a finding that motivates the targeting of serpentinizing systems for life detection efforts in our solar system, as well as further analysis of efforts to leverage serpentinites for industrial-scale geological H_2_ and carbon drawdown projects.

## Introduction

The vast majority of microorganisms exist in the Earth’s deep subsurface: oceanic sediments and the submarine and continental crust (1). It is thought that the bulk of microbial life in the subsurface exhibits biomass turnover on the order of years to thousands of years (2–4). Thus, the deep biosphere is thought to comprise the “slow majority” of microbial life on Earth (1, 3, 5). Life in the Earth’s crust faces substantial challenges, as the subsurface is often deprived of carbon and nutrient inputs from the surface and instead relies upon energy sources derived from radiolytic or water-rock reactions.

Serpentinization, the hydration and oxidation of ultramafic rocks, is a water-rock reaction that generates [hyper]alkaline, highly reducing fluids rich in H_2_, CH_4_, and low molecular weight organics (6, 7), electron donors that can support microbial activity. Despite living in the presence of abundant reducing power, life in serpentinizing systems faces numerous challenges including high pH (pH 8 – 12) and severe carbon/oxidant limitation (8–15) which may depress microbial metabolism and drive specific alkaliphilic adaptations (9, 16– 18). Thus, it remains unclear how the additive stressors of subsurface and serpentinizing conditions may affect the carbon assimilation rates and preferences of microorganisms – efforts to quantify such rates in serpentinizing systems have not yet been conducted.

Serpentinization of ultramafic mantle rocks has occurred through Earth history and is most often observed where Fe(II)-rich mantle rocks are uplifted along seafloor ridge systems or obducted onto land (19, 20). Serpentine has been identified on the surface of Mars and points to the possibility that aqueous alteration of ultramafic minerals occurred in the past and may still be occurring in the subsurface (21–23), where H_2_ gas could fuel chemosynthetic microbial ecosystems (21, 24). On Earth, interest in serpentinization has rapidly spread to the industrial energy sector. Due to the production of H_2_ during serpentinization, industrial development is underway to find deposits of natural H_2_ that have been preserved in the subsurface, or to stimulate geological H_2_ (GeoH_2_) through engineered water/rock reactions (25–28). Furthermore, serpentinizing systems have been targeted for *in situ* mineral carbonation, a carbon capture and sequestration (CCS) strategy wherein CO_2(aq)_ is injected into the subsurface to be stabilized as CaCO_3(s)_ (29–31). Both GeoH_2_ and CCS efforts could be negatively impacted by the microbial uptake of H_2_ or CO_2_, both of which may lead to fouling of engineered systems, consumption of hydrogen or the undesired biogenic conversion of CO_2_ into other byproducts than carbonate minerals, such as methane. It is necessary to determine the potential perturbations associated with microbial activity, even where life will be challenged by [hyper]alkalinity and carbon/oxidant limitation, to assess whether microbes will operate at the exceedingly slow rates so far observed in the subsurface biosphere.

Much of the prior microbiological work in serpentinizing systems has been conducted at fluid mixing interfaces, such as in marine systems, where hydrothermal effluent from serpentinizing rocks is released into oxidizing ocean water, or in continental surface-expressed seeps where discharging fluids are exposed to oxygenated atmosphere and surface-derived organic carbon (10, 20). These interfaces are energetically-rich, due to the strong redox disequilibrium between reducing serpentinized fluids and the oxidizing environments with which they mix. However, much lower energy availability, as well as carbon and nutrient abundances, are suspected in the bulk of the serpentinite rock-hosted subsurface, which may in-turn reduce microbial habitability and activity. Fortunately, it is possible to test this assumption with scientific drilling. We are able to access to the heart of the serpentinite-hosted biosphere within an actively serpentinizing ophiolite body via boreholes and rock cores generated by the Oman Drilling Project (ODP) and the International Continental Scientific Drilling Program (ICDP) in 2017-2018 at the Samail Ophiolite, the world’s largest body of terrestrial serpentinizing peridotite (19). This formation hosts aquifers with waters ranging from neutral to hyperalkaline (pH 7 – 12) and with a diversity of groundwater geochemistry (14, 32). Three continuous rock cores, BA1B, BA4A, and BA3A, were extracted, leaving behind observatory boreholes that provide access to serpentinite-hosted groundwaters. The BA1B, BA4A, and BA3A cores reveal a range of dunite and harzburgite lithology hosting a diversity of primary and secondary mineral components and exhibiting varying degrees of aqueous alteration (14, 19, 32, 33). Concurrently, the BA1B, BA4A, and BA3A boreholes host groundwaters exhibiting a gradient of aqueous geochemistry, ideal for studying the diversity of subsurface aqueous conditions and associated microbial activity that may exist (14).

Following recent work describing the abundance, structure, and genomic potential of microbial communities hosted in the Samail ophiolite, major questions remain as to the activity and carbon preferences of this rock-hosted microbiome. It is currently unknown which forms of carbon support microbial primary productivity in this system, given the [hyper]alkaline conditions and negligible inputs of organic carbon from the hyperarid surface regolith. It is also unclear to what extent the subsurface microbiome may respond to perturbations of H_2_ or DIC (27). Will physiological challenges of [hyper]alkaline, oxidant-limited, highly-reducing conditions limit microbial productivity; or is the microbial community capable of a response? To address these open questions, we constrain the rates and preferences of microbial carbon assimilation in the serpentinite-hosted subsurface through application of single-cell stable isotope probing (SIP) enabled by nanoscale secondary ion mass spectrometry (nanoSIMS). We probe the carbon sources that preferentially support biogenic methanogenesis by cavity-ringdown spectroscopy SIP (CRDS-SIP) of emitted CH_4_. With estimates of microbial biosynthesis, we apply a novel approach to estimate the catabolic rates of the subsurface microbiome and infer potential landscape-scale fluxes of H_2_ and CH_4_.

## Results and Discussion

### Accessing a gradient of subsurface serpentinizing groundwaters

We accessed groundwaters spanning the serpentinization reaction progression that leads to increases in pH and dissolved Ca, while decreasing DIC, dissolved Mg and Si (28), via the Oman Drilling Project Active Alteration Multi-Borehole Observatory. This borehole array consists of uncased 10 cm diameter boreholes 300 – 400 meters deep within a large body of serpentinizing mantle peridotite at the Samail Ophiolite, located within the Sultanate of Oman (14, 16, 29). The cores recovered from drilling reveal a serpentinized dunite/harzburgite-rich lithology containing relict primary minerals (e.g., olivine, pyroxene), diverse secondary minerals, and numerous generations of serpentine (14, 16, 29, 30). Core mineralogy varies at the reservoir scale and maps onto porewater geochemistry (14). Intact rock cores recovered from the drilling operation, as well as the aqueous geochemistry in the boreholes, have been previously described (14, 19, 32, 33). Groundwaters from the resulting boreholes have been monitored and pumped since initial drilling in 2017-2018 (14). For this study, discrete subsurface groundwater samples were accessed from three observatory boreholes at this site: BA3A, BA4A, and BA1B at three depths between 20 and 270 meters below land surface (mbls). Borehole BA3A contains hyperalkaline, Ca-OH type (pH 10.3 – 11.7), highly reacted, reducing fluids rich in H_2_, CH_4_, NH_3_, and Ca^2+^ (14). In contrast, borehole BA1B hosts mildly alkaline (pH 7.8 – 8.9), Si and Mg-rich groundwaters with greater oxidant availability and is denoted as an Mg-HCO_3_ type fluid. Borehole BA4A represents an intermediate of these two endmember fluids, containing moderately alkaline fluids (pH 9.2 – 9.9) of mixed composition. Key geochemical characteristics and cell abundances of these groundwaters are summarized in **Table 1**. For clarity, the waters hosted in BA1B, BA4A, and BA3A are hereafter referred to as “mildly alkaline,” “moderately alkaline,” and “hyperalkaline,” respectively.

**Table 1.** Abbreviated Summary of Groundwater Characteristics. Complete datasets for January 2023 borehole characteristics are reported in **Supplementary Text, Table S7, Supplementary Data.** All depths are reported as meters below land surface (mbls). DOC concentrations are displayed in **Fig. S11** and reported in **Supplementary Data**.

| Borehole | Description | Depth (mbls) | pH | EC (μS/cm) | Ca (mg/L) | Mg (mg/L) | Si (mg/L) | ΣS<sup>2-</sup> (μM) | Acetate (μM) | Formate (μM) | DIC (μM) | Cell Density (cell/mL) |
| --- | --- | --- | --- | --- | --- | --- | --- | --- | --- | --- | --- | --- |
| BA1B | Mildly Alkaline (Mg-HCO<sub>3</sub><sup>-</sup>) | 20 | 7.83 | 498 | 17.3 | 23.4 | 1.0 | 4.1 | 1098.02 | ND | 949 | 3.12×10<sup>6</sup> |
|  |  | 150 | 8.94 | 368 | 13.4 | 21.5 | 4.0 | 11.7 | 190.63 | 0.44 | 1733 | 1.12×10<sup>6</sup> |
|  |  | 250 | 8.08 | 491 | 22.1 | 30.0 | 6.9 | 79.7 | 443.74 | ND | 2385 | 3.88×10<sup>5</sup> |
| BA4A | Moderately Alkaline | 20 | 9.22 | 491 | 4.8 | 29.1 | 0.6 | 108.6 | 0.33 | ND | 1802 | 2.22×10<sup>6</sup> |
|  |  | 150 | 9.91 | 604 | 6.0 | 34.3 | 0.6 | 17.1 | 1.10 | 0.55 | 891 | 6.91×10<sup>5</sup> |
|  |  | 270 | 9.54 | 507 | 8.5 | 32.6 | 0.6 | 411.5 | 1.43 | 0.77 | 1685 | 8.38×10<sup>5</sup> |
| BA3A | Hyperalkaline (Ca-OH) | 20 | 10.32 | 1439 | 113.1 | 2.4 | 0.5 | 5.5 | 1.32 | 0.55 | 111 | 1.55×10<sup>6</sup> |
|  |  | 150 | 11.44 | 2960 | 180.5 | 1.0 | 0.5 | 5.2 | 1.76 | 0.88 | 48 | 6.92×10<sup>5</sup> |
|  |  | 270 | 11.72 | 3330 | 235.1 | 0.4 | 0.6 | 3.4 | 0.99 | 0.88 | 29 | 6.44×10<sup>4</sup> |

### Carbon assimilation preferences and rates of the serpentinite-hosted microbiome

We measured the carbon assimilation rates of 6323 individual microbial cells collected from the 9 different serpentinite sites (3 boreholes, each sampled at 3 depths) via single-cell stable isotope probing enabled by nanoscale secondary ion mass spectrometry (nanoSIMS-SIP). Anaerobically-sampled groundwaters from the boreholes described above were incubated with one of three ^13^C-labeled compounds: acetate (CH_3_^13^COO^-^), bicarbonate (H^13^CO_3_^−^), and formate (H^13^COO^-^) at near-ambient concentrations (**Supplementary Text, Table S1**). Incubations of mild- and moderately-alkaline groundwaters (BA1B, BA4A) were incubated for 40 days and those of hyperalkaline groundwaters (BA3A) were incubated for 238 - 254 days **(Supplementary Data)**. Incubations were conducted under H_2_ (2 atm) headspace ([H_2(aq)_] = 1.49 mM) (**Supplementary Text**) so as to provide groundwaters with sufficient H_2_ to sustain hydrogen-dependent metabolisms in the absence of continual H_2_ production from the host rock.

We measured the ^13^C enrichment of individual cells via nanoSIMS and calculated the cell-specific carbon assimilation rate, reported in fmol C cell^-1^ day^-1^, for each specific carbon tracer (34, 35). Across the groundwaters examined, we observe a range in assimilation rates between 10^-5^ and 10^-1^ fmol C cell^-1^ day^-1^ (**Fig. 1**). Carbon assimilation rates were significantly different between the boreholes (ANOVA, F_2, 6298_ = 763.41, p < 2.2e-16) with the slowest rates observed in hyperalkaline groundwaters (BA3A), where median carbon assimilation was 10^-5^ to 10^-3^ fmol C cell^-1^ day^-1^ (**Table S2**). In mild- and moderately-alkaline waters (BA1B, BA4A), microbial C assimilation was relatively uniform across depth, contrasted with hyperalkaline waters (BA3A). In hyperalkaline waters, appreciable microbial activity is detected only at shallow (20 m) depths (**Fig. 1, Table S3**) where 53 to 89% of cells exceed our activity threshold (cells whose lower error in ^13^C enrichment exceeds 3 SD than the mean ^13^C of negative control, see *Materials and Methods*). A large proportion of the incubated cells assimilated the ^13^C-probes across the mild- and moderately-alkaline waters (BA1B: 65 to 99%, BA4A: 50 to 100%) (**Table S3**). Below 150 m depth in hyperalkaline groundwaters (BA3A), only a fraction (0 – 37%, **Table S3**) of the microbial community exceeds our activity threshold, indicating that the majority of cells analyzed in these samples are dead, dormant, active only at exceedingly slow rates, or reliant on other, un-traced forms of carbon. Single-cell biomass, measured via nanoSIMS, appeared also to reflect groundwater and anabolic activity, with cells in hyperalkaline fluids being the smallest on average (BA3A: 53.6 fg C) followed by moderately alkaline fluids (BA4A: 71.9 fg C) and mildly alkaline fluids (BA1B: 90.0 fg C) (**Fig. S5**).

**Figure 1.**
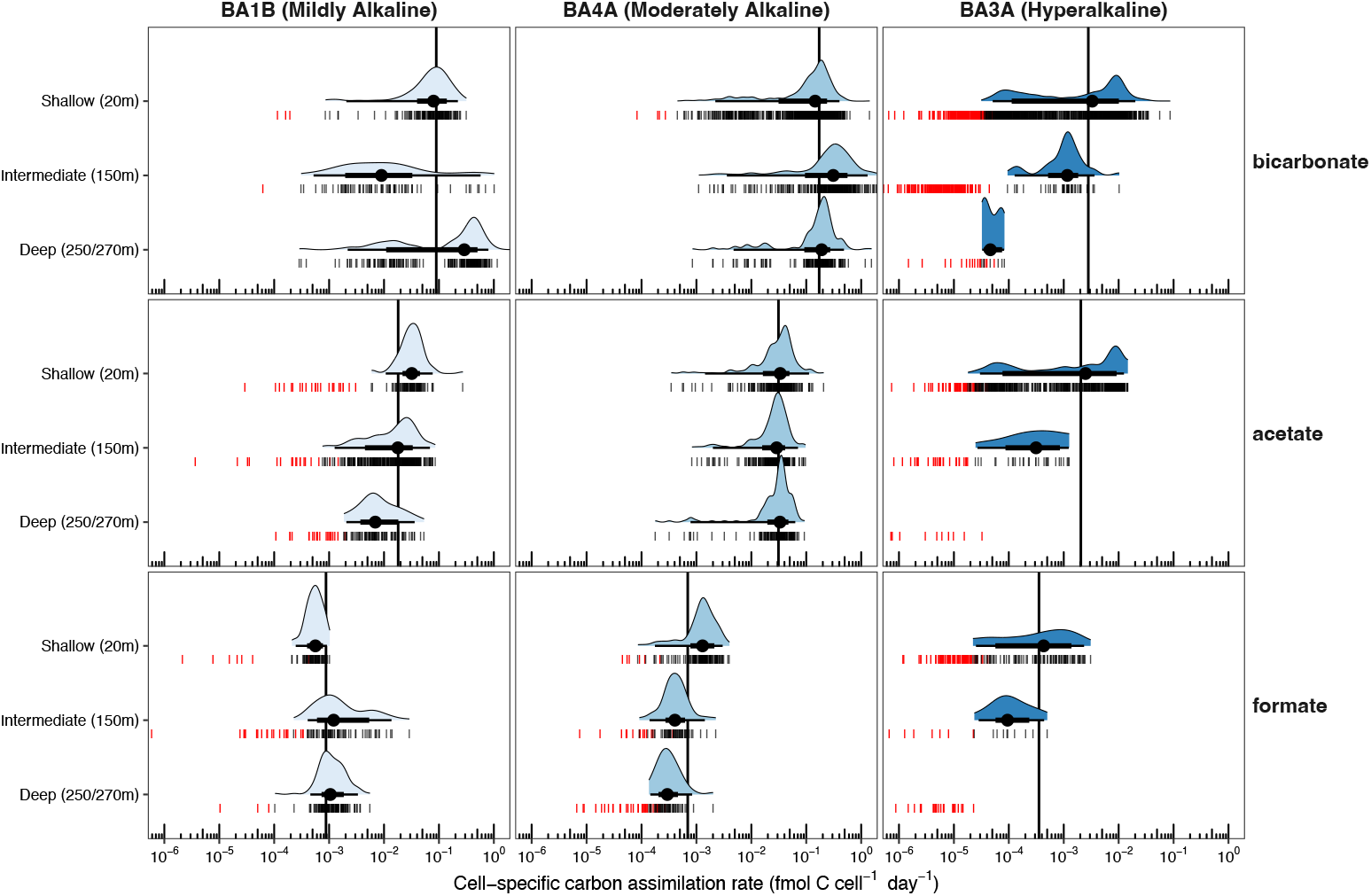
Cell-specific carbon assimilation rates across borehole, depth, and ^13^C tracer. For point-intervals: large points indicate median, bars represent 66% and 95% CI. Each point (vertical bar) indicates an individual single-cell measurement. Red points indicate measurements of ^13^C enrichment that did not exceed the cutoff for activity (Materials and Methods); these data are not included in calculations for point-intervals and density plots. Long vertical bars indicate the mean value within each panel (across depth). Note that BA4A and BA3A samples were acquired at 270 m; deep BA1B samples were acquired at 250 m.

Carbon assimilation rates were significantly different between the three carbon sources examined (ANOVA, F_2, 6296_ = 677.8, p < 2.2e-16), with bicarbonate being assimilated at the fastest rates, followed by acetate, then formate (**Fig. 1**). We had originally hypothesized that organisms hosted in hyperalkaline groundwaters (BA3A) would exhibit a distinct preference for low molecular weight organics (e.g., formate and acetate) given the exceedingly low ambient availability of inorganic carbon. However, ^13^C-bicarbonate is the only carbon tracer whose assimilation is detected in deep hyperalkaline fluids (4.61 × 10^-5^ fmol C cell^-1^ day^-1^) and bicarbonate is more readily incorporated in shallow and intermediate depth hyperalkaline fluids (10^-3^ fmol cell^-1^ day^-1^) relative to acetate and formate (9.5 × 10^-5^ to 2.5 × 10^-3^ fmol C cell^-1^ day^-1^). This observation is notable given that DIC becomes extremely limited in hyperalkaline fluids, which exhibit pH up to 11.7 and [DIC] as low as 29 µM (**Table 1**) suggesting that organisms in [hyper]alkaline groundwaters are particularly well-adapted to carry out carbon fixation under extreme DIC limitation (36). Formate is assimilated at the lowest rates across all depths and boreholes examined, in contrast with previous radiotracer measurements of formate oxidation in fluids from nearby sites (9), suggesting that formate may serve a primarily dissimilatory, rather than assimilatory role. We note that low apparent rates of ^13^C assimilation for any given cell do not necessarily imply a slow anabolic rate but may also indicate the shunting of ^13^C tracer through cross-feeding processes (i.e., oxidation of ^13^C-acetate or ^13^C-formate to ^13^C-DIC followed by fixation into biomass).

### Methanogenic archaea are pervasive and exhibit a strong preference for DIC

Methanogenic archaea, namely genus *Methanobacterium*, that transform hydrogen and dissolved inorganic carbon (i.e. CO_2_, HCO_3_^−^) to methane, are abundant members of the microbial community, including under hyperalkaline conditions. Assessed by 16S rRNA amplicon sequencing, the greatest relative abundances of *Methanobacterium* are observed in hyperalkaline groundwaters (BA3A: 29.5 - 45.2%) (**Fig. S3**), followed by moderately alkaline (BA4A: 12.0 - 18.5%), and mildly alkaline groundwaters (BA1B: 11.0 - 24.0%). Over the course of our incubations, *Methanobacterium* increased in relative abundance, from, on average 37% to 49% in hyperalkaline (BA3A), 15 to 55 % in moderately alkaline (BA4A), and 15 to 51% in mildly alkaline waters (BA1B) (**Fig. S3**) across all carbon sources tested, indicating a robust response to DIC and/or H_2_-rich conditions. *Methanobacterium* are identified alongside an array of nitrogen- and sulfur-cycling organisms observed previously in the Samail ophiolite as well as in other subsurface rock-hosted systems (**Supplementary Text**) (11, 37).

The nanoSIMS-SIP dataset described above is agnostic to taxonomy and likely captures a variety of organisms and metabolic strategies. To specifically quantify the activity and carbon preferences of methanogenic archaea, we applied cavity ringdown spectroscopy stable isotope probing (CRDS-SIP) (**Supplementary Text**) to measure the production of ^13^CH_4_ sourced from the SIP tracers. Methane production was consistent across mild and moderately alkaline waters (BA1B, BA4A: 10^-2^ µmol (mL day)^-1^) but was substantially lower in hyperalkaline waters (BA3A) (10^-3^ to 10^-4^ µmol (mL day)^-1^), with the deepest of these waters exhibiting barely detectable rates of methane production (**Fig. 2, Fig. S6**). We observed massive (> 75 at. % ^13^CH_4_) isotopic enrichment in samples incubated with ^13^C-bicarbonate (**Fig. 2)**. Serpentinite-hosted fluids can exhibit a range of DIC concentrations, from mM to sub µM, depending on the fluid type (38, 39). The groundwater recovered for this study exhibited DIC concentrations between 29 and 2300 µM, yet, even in shallow hyperalkaline fluids (BA3A, 20 m depth, pH = 10.32), where DIC = 111 µM, DIC accounted for > 75% of methane produced. The dominance of hydrogenotrophic methanogenesis in alkaline-to-hyperalkaline groundwaters indicates that oxidant and carbon limitation do not suppress methanogenic capacity for DIC utilization. Rather, methanogenic organisms are poised to produce CH_4_ from DIC across most of the conditions observed. High ^13^C enrichment of CH_4_ in shallow hyperalkaline fluids (BA3A, 20 m) suggests that deep, H_2_-rich hyperalkaline fluids, if perturbed and adequately relieved of oxidant and/or carbon limitation, may exhibit high rates of DIC-sourced methane production.

**Figure 2.**
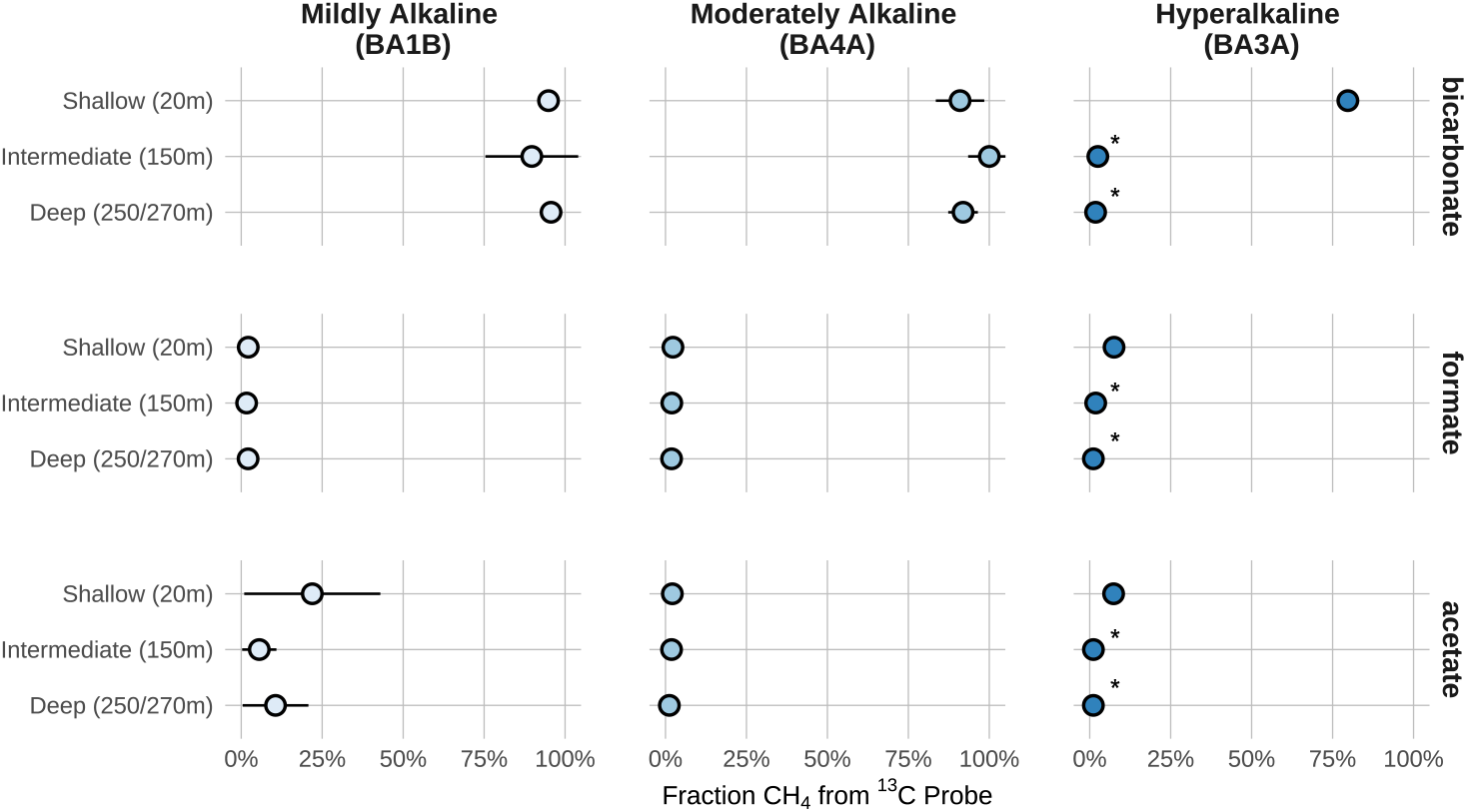
CRDS-SIP results. The fraction of biogenically-produced methane sourced from the ^13^C-tracer (^13^C-bicarbonate, ^13^C-formate, or 1C-^13^C-acetate) versus endogenous carbon sources is plotted. Concentrations of endogenous carbon sources, at natural carbon isotopic abundances, are reported in **Table S1**. Samples with total CH_4_ concentrations below detection by CRDS are marked with an asterisk (*) and should be considered null values. Raw ^13^F_CH4_ values are plotted in **Fig. S4**. Note that deep BA4A and BA3A samples were acquired at 270 m; deep BA1B samples were acquired at 250 m.

Our CRDS measurements reveal only minor contributions from formate to CH_4_ production. This result was surprising in light of metagenomic and radiotracer work indicating the capacity for formatotrophic methanogenesis in the Samail ophiolite (9, 40, 41). We observe the highest contribution of ^13^C-formate to produced methane, 7.5%, in shallow hyperalkaline fluids (BA3A), supporting previous work indicating that formatotrophy may be an adaptation to hyperalkaline waters (9). However, our finding that DIC is consistently preferred over formate contrasts with prior work that was unable to measure DIC-sourced CH_4_ production in hyperalkaline fluids at a nearby hyperalkaline borehole (NSHQ14) (9). This may be due to differences in measurement technique (CRDS vs. radiotracer), and incubation length (36 weeks vs. 8 weeks).

Mildly alkaline groundwaters (BA1B) incubated with 1C-^13^C-acetate reveal 5.6 to 21.9% of sourced methane sourced from the ^13^C label. We considered two hypotheses to explain this result: (i) acetoclastic methanogens are present or (ii) acetate oxidation to DIC supports hydrogenotrophic methanogenesis through direct or indirect syntrophy. Despite energetic favorability (38), the genomic capacity for acetoclastic methanogenesis by *Methanobacterium* in has yet to be identified in subsurface Samail ophiolite fluids (9, 41, 42). Because our acetate ^13^C probe was labeled at the C1 position, the acetoclastic pathway would result in labeling of biomass but not CH_4_, as methane is sourced from the C2 (methyl) component of acetate (43). The presence of ^13^CH_4_ production from a 1C-^13^C-acetate label demonstrates that acetate must have been oxidized to DIC prior to its entry into a methanogenic pathway. Therefore, we find the hypothesis of syntrophic acetate oxidation (ii), either direct or indirect, to be more plausible. While the origin of abundant acetate in mildly alkaline groundwaters (BA1B, **Table 1**) remains unclear, our results indicate that it is likely oxidized to DIC (versus direct conversion to methane) prior to its subsequent remobilization as CH_4_.

### Potential bioenergetic fluxes suggest large biological contribution to subsurface geochemistry

The apparent dominance of DIC as a carbon source for both anabolic and methanogenic activity under [hyper]alkaline conditions motivates an exploration of the potential magnitude of microbial activity on the reservoir scale. To this end, we apply single-cell C assimilation rates to estimate the bioenergetic flux of the serpentinite-hosted microbiome by calculating biological mass-specific power (MSP) (44, 45). Metabolic processes and microbial activity in natural systems may be poorly predicted by estimates of free energy availability because free energy calculations rely on measurements of metabolites in standing pools rather than their rates of transformation (46). By deriving bioenergetic flux from empirically-measured rates of C assimilation, we sidestep difficulties with estimating metabolic potential from chemical disequilibrium. We posit that directly-measured carbon assimilation rates offer insight into the energy throughput of subsurface microorganisms. The production of cell biomass, the anabolic rate (mass C per unit time), directly relates to the power (energy per unit time) generated by metabolism (44, 45). Therefore, we calculate MSP as watts per gram of biomass (W (g C)^-1^) with an empirically-derived relationship of power requirements and cellular biosynthesis (**Materials and Methods**) (45). MSP provides a common basis for comparing bioenergetic flux across a broad range of metabolisms and ecologies, and for evaluating organismal energy flux relative to basal maintenance costs.

MSPs inferred from our ^13^C-SIP experiments are notably high (**Fig. 3**). Averaged across all carbon sources examined, we observed the highest community-level average MSP of 9.4×10^-3^ W (g C)^-1^ in shallow alkaline fluids (BA4A 20 m) and the minimum community-level average MSP of 8.5×10^-6^ W (g C)^-1^ in deep hyperalkaline fluids (BA3A 270 m). Lower bounds on MSP are proposed by Lever et al. (47) as the temperature-dependent energy costs of amino acid racemization repair, and such limits are on the order of < 10^-6^ W (g C)^-1^ (**Fig. 3**). Therefore, we infer that the majority of cells sampled in this study are operating with energetic fluxes orders of magnitude greater than basal maintenance costs that might be expected for organisms in dark, oligotrophic, [hyper]alkaline, subsurface aquifers. Because ^13^C-labeled substrates are not passive tracers of cellular biosynthesis, ^13^C assimilation rates (and by extension, MSP) may fairly be interpreted as lower bounds on cellular biosynthesis (**Eq. 5**), as cells may be acquiring carbon from non-traced sources. While lower MSPs we observe are akin to organisms inhabiting deep marine sediments, these values may likely reflect cells that are actively assimilating non-labeled carbon sources or are assimilating ^13^C through syntrophic cross-feeding reactions, a process we infer to be occurring based on our CRDS-SIP measurements, described above. However, the fact that the majority of the microbial community exhibits a high apparent MSP in serpentinizing conditions is notable. The microbial communities hosted in alkaline groundwaters exhibit median MSP equivalent to microbiomes in surface or near-surface soils and shallow aquatic sediments (**Table S4**). Similarly, the upper MSP we observe (0.04 W (g C)^-1^) is approximately that of the average human being (0.012 W (g C)^-1^), which hardly conforms to the notion of the ‘starving majority’ of microbes in subsurface environments (**Fig. 3**) (45). This result is surprising in light of previous measurements of microbial activity in both terrestrial and marine subsurface environments that imply the ubiquity of low activity (2). Our estimates of MSP suggest that microbes in serpentinite-hosted groundwater are markedly capable of high bioenergetic flux, especially under perturbed conditions.

**Figure 3.**
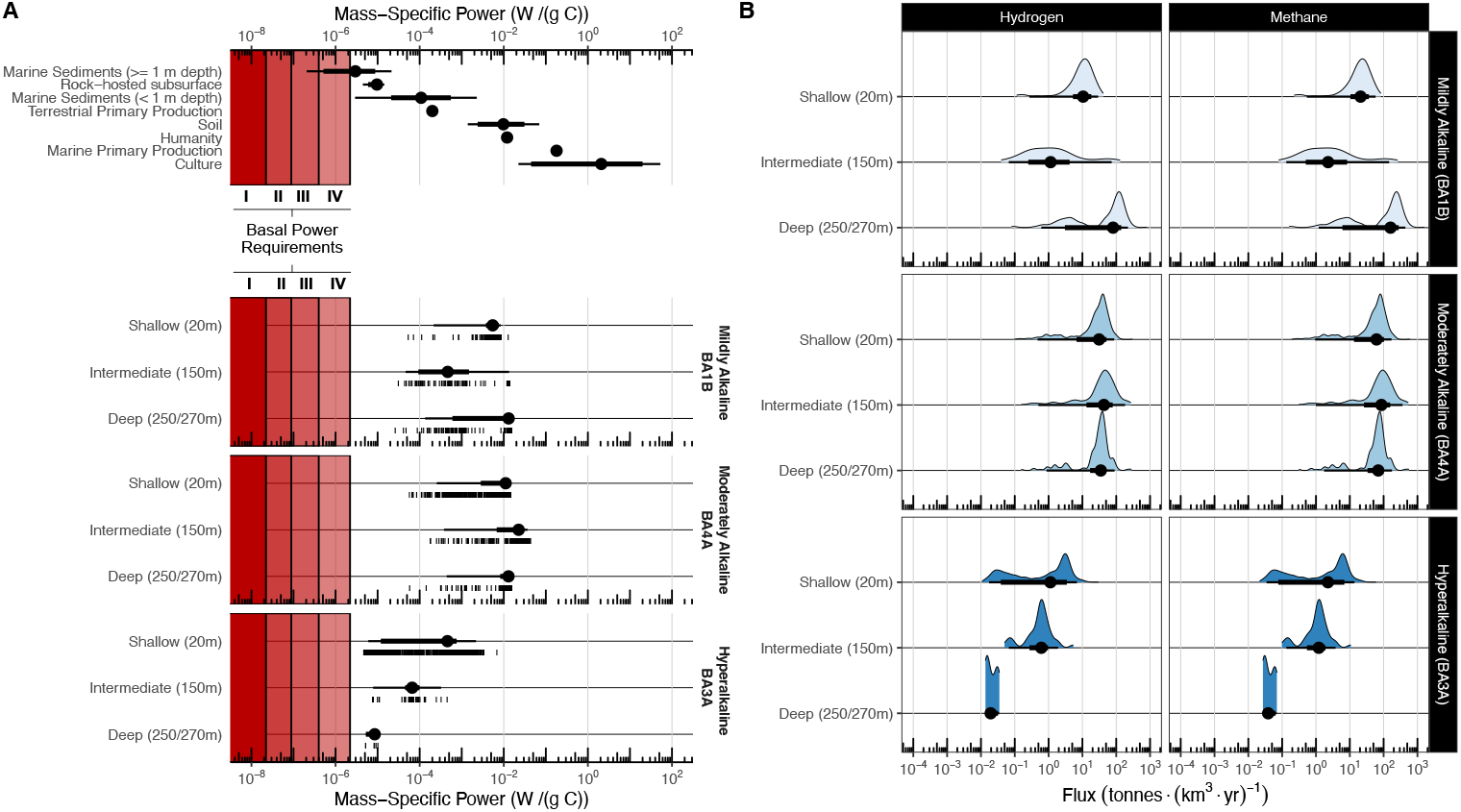
**(A)** Top panel: literature-derived estimates of biological mass-specific power from a variety of Earth systems (45). Bottom panel: estimates of mass-specific power derived from our single-cell ^13^C assimilation rate measurements. Only cells with cell-specific carbon assimilation above quantification limits are plotted (**Materials and Methods**). Basal power requirements are defined as the minimum energy required to repair spontaneous amino acid racemization at 35^°^C. Four scenarios, postulated by Lever et al. (47), are annotated by red vertical bars in order of increasing energetic cost: (i) single amino acid excision and replacement (MSP > 2×10^-8^ W (g C)^-1^), (ii) protein replacement with 2% racemization, (iii) protein replacement with 10% racemization, and (iv) full protein replacement (MSP > 2×10^-6^ W (g C)^-1^). **(B)** Ranges of estimated H_2_ oxidation and CH_4_ production rates are plotted. For these estimates, only hydrogenotrophic methanogenesis is considered. For point-intervals in all panels, large point indicates median, thick and thin bars represent 66% and 95% CI, respectively.

MSP is a useful metric to calculate because it enables us to estimate the potential fluxes of cryptically-cycled metabolites. Here we specifically focus on estimating biotic H_2_ oxidation and CH_4_ production, as these fluxes are relevant to future geoengineering efforts. The net H_2_ production of peridotite in the Samail ophiolite is difficult to estimate due to uncertainties in both gross abiotic production and biological oxidation processes. H_2_ fluxes measured at the surface integrate both production and consumption terms (27, 48). Similarly, because CH_4_ has both biotic and abiotic sources, rates of biological methanogenesis are difficult to predict at the landscape scale, though isotope geochemical approaches have indicated a potentially large contribution (36, 38). The predicted microbial production of CH_4_ and the consumption of H_2_ can be estimated at the reservoir scale by applying MSP estimates to well-constrained ecosystem parameters. We first convert MSP into volume-specific power (VSP: the sum flux of metabolic energy in a volume of rock). We then calculate annual H_2_ oxidation and CH_4_ production per cubic kilometer of rock by multiplying VSP by the molar yield of the metabolism, assuming an actively serpentinizing layer of 1 km depth (49). These rates can be of substantial magnitude. Median estimates for microbial H_2_ consumption vary between 0.02 and 80.6 tonnes H_2_/km^3^/yr, with greater consumption in mild and moderately alkaline fluids (BA1B, BA4A) and lowest consumption in hyperalkaline fluids (BA3A) (**Fig. 3, Table S5**). Calculated H_2_ consumption rates are in excess of or on par with measured rates of H_2_ efflux from the Samail ophiolite: 2 - 4 tonnes H_2_/km^3^/yr (27, 50), indicating that gross abiotic H_2_ production may be far greater than would be suggested by efflux of H_2_ from the system which, instead, represents the net of production and consumption. Such rates of microbial H_2_ oxidation also suggests that a majority of H_2_ produced by low-temperature serpentinization reactions is removed in mild and moderately alkaline fluids and that accumulations of H_2_ in hyperalkaline fluids may be primarily the result of depressed microbial consumption.

Similarly, we estimate median CH_4_ emission factors via MSP to be between 0.04 to 160 tonnes CH_4_/km^2^/yr (**Fig. 3, Table S5**), with similar trends across serpentinizing groundwater chemistry. These fluxes may be significant at the landscape scale. In rocks hosting mild- and moderately-alkaline groundwaters, these emission factors are comparable to those defined by the IPCC for agricultural and brackish ponds (∼10^1^ tonnes CH_4_/km^2^/yr) and enteric fermentation emissions from cattle (∼10^1^ tonnes CH_4_/km^2^/yr) (51, 52). We note that we report CH_4_ emission factors in units of surface area rather than volume, assuming a reaction depth of 1km, such that our fluxes are comparable to those as reported by the IPCC. We further note that our estimated fluxes carry major uncertainties regarding the subsurface geological and biological heterogeneity and the extent of trapping vs. emission of gases but emphasize that our purpose in conducting these estimates is to provide wide constraints on biological contributions to the system at scale. Such magnitudes of observed biological activity may result in non-trivial conversions of hydrogen and CO_2_ to methane at rates that could significantly impact the utility of industrial-scale GeoH_2_ and CCS efforts within ultramafic rocks. Furthermore, these estimates provide useful means of comparison to other Earth systems where microbial contributions to landscape or global biogeochemistry have been modeled (35).

### Implications of a robust serpentinite-hosted microbiome

Serpentinizing environments are unique for hosting H_2_-rich groundwaters that challenge microorganisms with carbon, oxidant, and nutrient limitation under reducing, [hyper]alkaline conditions. The forms of carbon that support the subsurface microbiome in these environments have remained elusive but are important to understand in order to generate predictions of subsurface habitability and response. The results of our anabolic (^13^C-nanoSIMS-SIP) and methanogenic (CRDS-SIP) tracer experiments indicate DIC as the preferred substrate for both assimilatory and methanogenic processes across all groundwaters tested. Rates of carbon assimilation vary widely between the different carbon sources tested, indicating stark differences in carbon assimilation rates among the taxonomic groups and groundwater chemistries identified herein. The anabolic activity captured with our nanoSIMS-SIP dataset may reflect the activities of a wide-variety of autotrophic, formatotrophic, and acetotrophic organisms. For example, while *Methanobacterium* is abundant in these groundwaters, autotrophs such as acetogens are also abundant in Samail ophiolite groundwaters and may contribute to C fixation (11, 16, 39, 53). CRDS-SIP and 16S amplicon sequencing, however, provide strong evidence that methanogens are highly active members of the subsurface community and exhibit a distinct preference for DIC, even under [hyper]alkaline conditions. Carbon assimilation rates are highly variable across the groundwaters tested, exhibiting sharp drop-offs in hyperalkaline (BA3A) groundwaters where oxidant limitation becomes extreme. These conditions are likely inextricably tied to the hydrologic recharge characteristics of the host-rock, where reduced hydrologic connectivity results in decreased oxidant and carbon availability (**Fig. S12**) (54, 55).

Inherent to any SIP experiment are the uncertainties associated with experimental manipulation and perturbation. Thus, it is necessary to assess the degree to which our measured anabolic and methanogenic rates represent those in ambient *in situ* conditions. As described, our incubations were initiated at aqueous H_2_ concentrations (∼1.49 mM) that are typically above ambient concentrations for mildly-alkaline groundwaters (BA1B, 0.175 µM) and are more comparable to concentrations expected in moderately-alkaline and hyperalkaline fluids (BA4A, BA3A, 1-4 mM) (**Table S6**). Because half saturation constants (*K*_*s*_) for hydrogenotrophic archaea and bacteria are expected to be on the order of 10^0^ - 10^1^ µM (56) (though these values are unconstrained for organisms isolated from serpentinizing environments), supplying H_2_ in mildly-alkaline fluids (e.g., BA1B) could ostensibly support hydrogenotrophic growth at rates approximately 5X – 50X that of ambient rates in temperate fluids (BA1B), as estimated by Monod kinetics. Anabolic and methanogenic rates, measured here, as well as our inferred landscape-scale fluxes, could fairly be considered potential upper-bounds under H_2_-stimulated conditions, rather than true *in situ* rates, though organisms already saturated with H_2_ may not experience significant change. Even so, the exact degree of stimulation remains difficult to pinpoint. Variation in ambient H_2_ concentrations in Samail ophiolite groundwaters have been measured and exhibit differences based on borehole, depth, and sampling technique (**Table S6**). Furthermore, H_2_ appears to exhibit sporadic release events from peridotite fracture systems, suggesting that dissolved H_2_ may be spatially and temporally heterogenous (57). The method of experimental H_2_ introduction is also important to consider. Incubation vials experience a high initial pulse of H_2_ maintained in the headspace, whereas natural groundwaters receive sustained H_2_ flux from the host rock. Perturbed hydrologic mixing regimes, such as rock fracturing or fluid injection, may lead to elevated, sustained fluxes of H_2_ into groundwaters that are orders of magnitude greater than those observed in the native system due to elevated water-rock ratios (27). The application of ^13^C tracers may also have a stimulating effect. We consider potential stimulation by tracers to be less probable than H_2_ given that incubations with a greater degree of nutrient amendment over background were not those which exhibited the greatest anabolic rates (**Table S1**). Regardless of the exact degree of stimulation our methodology may have introduced, our main finding holds: the serpentinizing subsurface hosts a robust microbiome that is poised to consume H_2_ and DIC.

Our estimates of anabolic and energetic capacity in subsurface serpentinite-hosted organisms raise urgent questions for industrial efforts to conduct fluid injections into subsurface igneous rock formations. The mineral carbonation CCS process injects CO_2(aq)_ into the subsurface where it is immobilized as mineral carbonates (29, 30, 58). Microbial growth and activity can threaten CCS efforts, both through biofouling of hydraulic equipment, as was the case with CarbFix (59), or through the geochemical impact of microbial activity itself via CO_2_ consumption and production of unwanted byproducts including H_2_S and CH_4_. Similarly, GeoH_2_ production and recovery efforts may be compromised by undesired microbial H_2_ consumption that reduces net yields (27). The general perception of subsurface environments as starved ecosystems hosting slow-acting microbial life can lead to the erroneous assumption that the subsurface is amenable to industrial-scale perturbation with low risk. Rather, this work demonstrates that, across [hyper]alkaline groundwaters and partially-serpentinized bedrock, there exists an abundant, highly active microbiome whose constituents are poised to assimilate DIC and to use H_2_ to transform to CH_4,_ even at hyperalkaline pH. H_2_ produced by geological stimulation may therefore be rapidly consumed by microbial communities, especially those in mildly- and moderately-alkaline groundwaters. Similarly, injection of CO_2_-rich fluids into deep, hyperalkaline fluids (e.g., BA3A), may serve to decrease pH and increase [DIC]. If hyperalkaline fluids were altered in this way, their constituent microbial activity may rapidly increase towards rates observed in more mild/moderately alkaline waters (an increase of two orders of magnitude). Targeted follow-up studies are urgently needed to evaluate how organismal- and community-level changes driven by fluid injection into mafic and ultramafic rocks can affect reservoir-scale changes in H_2_ and CH_4_ dynamics. Specifically, characterization of how microbial activity is modulated by *in-situ* geochemical conditions will enable predictions of how these ultramafic systems will respond to engineered perturbations.

This study highlights serpentinizing systems as an important contrast to many other deep-subsurface or rock-hosted ecosystems. Most previous sites of deep biosphere research, such as deep marine sediments and deep granitic crust, are extremely energy limited in nature (1–3, 5, 35). Therefore, previous estimates of anabolic rates in the deep biosphere, despite being based on a variety of techniques and assays, have consistently inferred slow rates of microbial growth and activity, supported by trace energy derived from rock through radiolytic or water-rock reactions, or sedimentary organic matter (2). This work supports the notion that the redox gradients and dynamic hydrology produced by serpentinization processes provide ample energy for microbial activity in the absence of sunlight and surface-derived organic carbon (18, 60). We demonstrate that the serpentinite-hosted microbiome is poised for robust anabolic and methanogenic activity whose bioenergetic capacity rivals that of surface environments. Serpentinites on Earth or elsewhere in the solar system may therefore exhibit high capacities for microbial activity which improve the probability for the recovery of organic and chemical biosignatures. The recent discovery of aqueously altered igneous rocks and their association with organic matter by the Perseverance rover, as well as the proposed existence of liquid Martian groundwater hosted in an igneous mid-crust (23), underscores the relevance of serpentinization products as life detection targets on extraterrestrial bodies (22, 61, 62).

## Materials and Methods

### Groundwater sampling

Subsurface groundwater was accessed through boreholes BA1B, BA4A, and BA3A, located in the Oman Active Alteration Multi Borehole Observatory established by the Oman drilling project in 2018 (14, 19). Water samples, 970 mL each, were collected from discrete intervals using a point-source bailer connected to a distance-marked tag-line (Solinst). Groundwater was directly sampled from the bailer with a modified sample retrieval tool (Solinst) connected in-line with a sterile 60 mL syringe, which was used to aseptically distribute samples. For SIP-incubations, 50 mL of site fluid was injected directly into evacuated vials hermetically sealed with butyl stoppers. Vials were transported back to the lab at ambient temperature. For DNA sequencing, 100 – 300 mL of site fluid was filtered through 0.1µm PVDF filters. Filters were placed in 2 mL of DNA/RNA Shield (Zymo) and stored at −70^°^C in a liquid N_2_ dewar. Filtrate from the DNA samples was saved for IC, ICP-OES, and IRMS analyses, described below. Water samples for NMR analysis were filtered through 0.22µm PES filters and stored in combusted (450^°^C, 8 hr) amber glass VOA vials whose PTFE liners were triple-rinsed with methanol, dichloromethane, and hexane in series. pH, temperature, and conductivity were measured on-site with a handheld probe. Contamination of groundwater was assessed by inspecting samples for fluorescent pigment tracers injected in drilling fluid during initial coring operations (14). Fluorescent tracers, initially amended during drilling at an average concentration of 10^8^ particles/mL, were not detected in any samples.

### SIP Incubation

Prior to SIP incubation, the headspace of each vial was flushed with H_2_ and pressurized to 2 atm in order to supply sufficient H_2_ in the absence of H_2_ generated by serpentinization. By Henry’s law (adjusted for the temperature, 35^°^C, experienced by groundwaters in the system) (**Table S6**), this results in a dissolved H_2_ concentration of 1.49 mM H_2_. Samples were anaerobically amended with one of the following ^13^C-enriched tracers: sodium formate (Cambridge Isotope Laboratories Inc., CLM-583-1, ^13^C, 99%), sodium bicarbonate (Cambridge Isotope Laboratories Inc., CLM-441-5, ^13^C, 99%), or sodium acetate (Cambridge Isotope Laboratories Inc., CLM-156-5, 1-^13^C, 99%). Stocks of sodium acetate and sodium formate ^13^C-probes were prepared at concentrations of 5000 µM with 0.5mL injected (1:100 dilution). Sodium bicarbonate stocks were prepared as 600 mM, 100 mM, and 10 mM for BA1B, BA4A, and BA3A, respectively, with 0.5 mL injected for each (1:100 dilution). The resulting effective concentrations and isotopic composition stable isotope probes are reported in **Table S1**.

Groundwater samples were incubated at 35^°^C, analogous to the typical subsurface fluid temperature measured in the Samail Ophiolite, **(Table S6)** in a forced air incubator. Microbial activity was monitored by periodic measurement of headspace methane (Materials and Methods: Gas Measurements) and aqueous total sulfide. Total aqueous sulfide was measured by colorimetric (diamine) assay. These measurements informed the SIP incubation duration. For detailed quantification of sulfate reduction rates in this system, we refer the readers to prior work by Glombitza et al. (16). BA1B and BA4A samples were incubated for 40 days, BA3A samples were incubated for 238 - 254 days (**Table S8**).

At the end of the incubation period, 1mL of incubation fluid was anaerobically removed from incubation experiments with an N_2_-flushed needle and fixed in paraformaldehyde to a final concentration of 2%. Cells were isolated by centrifugation, washed 3X in MQ water, then resuspended in 100 µL of MQ water. 20 µL of cell suspension was spotted onto combusted (450^°^C, 8hr) aluminum-coated glass slides and allowed to air-dry before analysis. Aluminum coated glass slides were used as the sample substrate because, in contrast to silicon wafers often used for nanoSIMS analyses, aluminum does not exhibit a Raman signal that overlaps with cell spectra.

### Raman microspectroscopy

To identify microbial cells, but avoiding isotopic dilution effects of nucleic acid staining on ^13^C/^12^C measurements conducted via nanoSIMS (63), we applied single-cell Raman microspectroscopy to confirm the biogenicity of cell-like particles prior to analysis via nanoSIMS. Raman microspectroscopy was conducted at the University of Colorado Boulder Raman Microspectroscopy Lab using a Horiba LabRAM HR Evolution Raman spectrometer equipped with a 100 mW 532nm (green) excitation laser. The laser was focused with a 100X (NA = 0.90) air objective lens, resulting in a laser spot size of ∼1 µm. Single-cell spectra were captured in the 200 – 3150 cm^-1^ range using 100% laser power (2.55 mW) over one acquisition of 45 seconds. A 100 µm confocal pinhole and 600 lines/mm diffraction grating were used, resulting in a spectral resolution of ∼ 4.5 cm^-1^. Biogenicity was confirmed by inspection of raw spectra for diagnostic organic bands in the fingerprint region (200 – 1800 cm^-1^) as well as signature C-H stretching band at 2800 – 3100 cm^-1^ associated with lipids, proteins, and nucleic acids. Because Raman spectroscopy is a non-destructive technique, particles that were identified as cells were targeted for correlative nanoSIMS imaging. Images taken on the instrument with a CCD camera were used as guides for nanoSIMS imaging.

### Nanoscale secondary ion mass spectrometry

Samples were analyzed at the California Institute of Technology Microanalysis center with a CAMECA NanoSIMS 50L (CAMECA, Gennevilliers, France). Before analysis, samples were sputter-coated with a 25 nm layer of Au. Analysis chamber vacuum was held below 5×10^-8^ Torr for the entirety of the run. NanoSIMS was conducted with a 10 pA primary Cs^+^ beam current. Before acquisition, a presputter of 7 minutes with a 250 pA primary beam current was applied, and the instrument was tuned to achieve MRP > 5000. All frames were acquired with a 512 × 512 pixel raster covering a 25 × 25 µm area. For each area, 5 repeated frames were collected. The following masses were collected in parallel with electron multipliers: ^12^C^-, 13^C^-, 12^C^14^N^-, 18^O^-, 28^Si^-, 31^P^-, 32^S^-^. Isotopic enrichment of cells is reported relative to cells in groundwater samples without a ^13^C probe added. Single-cell isotope values were calculated using a custom script based on the *sims* Python library. Cell areas were manually defined using the GNU Image Manipulation Program (**Fig. S7**). Isotope ratios were converted to fractional abundances ^13^F using the relationship ^*13*^*F =* ^*13*^*R /(1 +* ^*13*^*R)* and ^*13*^*R =* ^*13*^*C /* ^*12*^*C* where ^13^C and ^12^C represent the total ion counts measured for each cell, accumulated over the 5 repeated acquisitions. A deadtime correction was also applied:

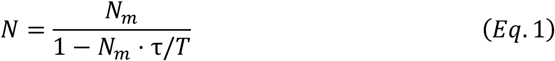

where *N* is the real number of secondary ions, *N*_*m*_ is the number of secondary ions measured, τ is deadtime (in seconds) and *T* is dwell time (seconds per pixel). The deadtime of the detectors on the instrument used for this study is 44 ns; the dwell time applied was 1600 µs.

### SIP calculations

Calculation of cell-specific carbon assimilation is carried out similar to the approach defined by Stryhanyuk et al. and Polerecky et al. (34, 64), which requires end-point measurement of isotopic enrichment after incubation of cells in the presence of an isotopically enriched tracer. *K* represents the source-normalized excess ^13^C atom fraction incorporated by a cell in the presence of ^13^C-labeled compounds. In other words, *K* represents the fraction of cellular carbon derived from a ^13^C tracer and is defined as:

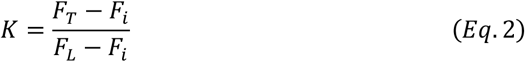

where F represents the ^13^C isotopic fractional abundance of biomass time of sampling (F_T_), biomass at the start of the incubation (F_i_), and the effective isotopic label strength (F_L_). To calculate cell-specific C assimilation rate, we convert estimated cell-specific volume to cell-specific mass using the relationship described in Khachikyan et al. (65): m_carbon_ = 197 × V^0.46^ where V refers to cellular volume in µm^3^ and m_carbon_ refers to cell-specific carbon mass in fg. Cell specific volume is derived from nanoSIMS ROI area by calculating the volume of two capping hemispheres on a coccoidal or rod-shaped cell (34, 35).

We consider two methods to derive the cell-specific carbon assimilation rate (r_C_). The first is similar to that outlined by Stryhanyuk et al. (34), where the cell-specific mass of carbon (*m*_*i*_) is multiplied by the source-normalized ^13^C atom fraction excess (K) per unit time (t):

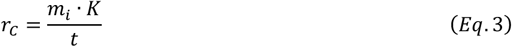

The second method is defined by Polerecky et al. (64), wherein the possibility of cell division occurring is taken into account:

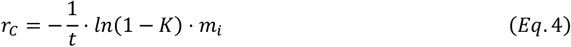

Because our 16S amplicon data suggests that community shifts have occurred, we report r_C_ values derived from Eq. 4. The number of cell divisions an individual cell has undergone is not possible to constrain with current methods, though we report differences between the two calculation methods that are minor (**Fig. S8**). We note that this work reports *r*_*C*_ in units of fmol C cell^-1^ day^-1^. Intracommunity heterogeneity in carbon assimilation is quantified using the Gini coefficient (**Supplementary Text, Fig. S1**).

A negative control was included to establish natural abundance ^13^C isotope ratios of the cells. Cells in SIP incubations whose isotopic enrichment or lower bound on propagated error did not surpass 3 standard deviations of the negative control, were considered “inactive” and excluded from our analyses and statistical tests. In other words, the limit of quantification is achieved when *F*_*cell*_ − σ_*cell*_ > F_CTL_ + (3 × *SD*_*CTL*_) where *F*_*cell*_ is the mean isotopic fractional abundance of a given cell region-of-interest, *σ*_*cell*_ is the propagated error of the isotopic fractional abundance, *F*_*CTL*_ is the mean isotopic fractional abundance of cells in the negative control, and *SD*_*CTL*_ is the standard deviation of isotopic fractional abundance in the negative control (**Fig. S10**).

### Calculations of MSP and metabolic flux

Heijnen & van Dijken established an empirical relationship between energy dissipation and biomass yield for organisms across a wide array of cultured organisms and substrates, finding that the average biomass yield when expressed in energy-based terms was 0.019 ± 0.008 and 0.03 ± 017 g C (kJ)^-1^ for aerobes and anaerobes, respectively (44). Data compiled from an array of environmental systems demonstrates that this relationship can be extended to much lower biomass yield rates such as those observed in environmental systems, indicating that energy-to-biomass conversion factors hold across Earth environments (45). The stability of biomass yield over orders of magnitude metabolic rate enables prediction of biomass-specific energetic flux. We consider the mass specific carbon turnover rate (µ*) in mass carbon biomass synthesized (C_biosynthesis_) per unit time per mass of standing biomass carbon (C_biomass_), or µ* = ((C_biosynthesis_) × s^-1^ × (C_biomass_)^- 1^). Mass specific power (MSP) is related to the biomass yield (Y* = C_biosynthesis_ /J) of an organism by:

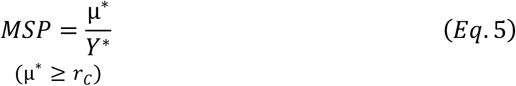

We note that, for the purposes of MSP calculations, cell-specific carbon assimilation (*r*_*C*_) as derived from nanoSIMS-SIP may be an under-estimate of aggregate biomass turnover, as cells may source biomass carbon from non-13C-labeled sources (e.g., syntrophic cross-feeding, or secondary metabolic pathways). Therefore, MSP values could fairly be considered lower bounds.

Biomass yield (Y*) for anaerobic organisms is related to the mass-specific carbon turnover rate (yr^-1^) by the linear relationship:

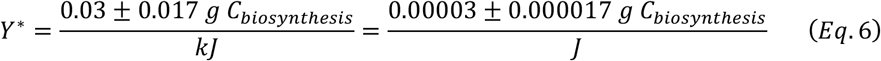

NanoSIMS-derived estimates of single-cell biomass turnover (µ*, fg C_biomass_ s^-1^), as measured in this study, are applied to estimate MSP (note that the units for each term are displayed in brackets for clarity):

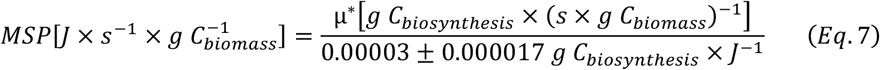

MSP is then converted into cell-specific power (CSP) via cell-specific biomass inferred from nanoSIMS measurements:

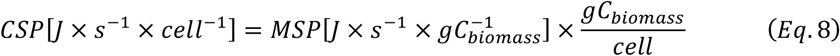

We then calculate an energy requirement in joules per unit volume of rock (volume-specific power, VSP) by multiplying CSP by a conservative rock-hosted cell density factor (average of 10^5^ cells/cm^3^) (14) and adjusting by the fractional abundance (*a*) of *Methanobacterium* (*0 < a < 1*):

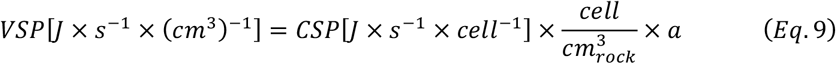

We then apply an estimate of 1.255 × 10^10^ Joules produced per tonne of H_2_ consumed in methanogenesis, a value derived from estimated Gibbs free energy (ΔG) yields of hydrogenotrophic methanogenesis, 4H_2_ + CO_2_ → CH_4_ + 2H_2_O, in Samail ophiolite waters (66). We calculate volume-specific H_2_ oxidation rate as:

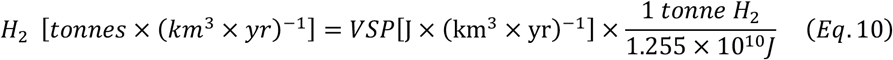

Because more favorable ΔG leads to a reduced catabolic flux, we also consider the Gibbs free energy yield under far-from equilibrium conditions (ΔG^°^), in this case 2.393 × 10^10^ Joules produced per tonne H_2_ (48.1 kJ/mol H_2_) as a lower constraint on catabolite flux (49) (**Fig. S9**).

In both cases, CH_4_ production is subsequently estimated by the stoichiometry of hydrogenotrophic methanogenesis.

We compare CH_4_ production (tonnes (km^3^ yr)^-1^ to IPCC-defined emission factors (51, 52) with the assumption that methane produced in a volume of serpentinizing rock 1 km in depth (1 km^3^) will efflux from a corresponding surface area (1 km^2^). IPCC emission factors for enteric fermentation by cattle (reported as kg CH_4_ per head) are converted to tonnes CH_4_ (km^2^ yr)^-1^ by applying a standard stocking density of 0.35 head/acre.

### Dissolved anion and cation analysis of site fluids

Water samples and control sample concentrations were analyzed for total concentrations of metals and major inorganic cations (including Al, As, B, Ba, Ca, Cd, Cu, Cr, Fe, K, Li, Mg, Mn, Mo, Na, Ni, P, Pb, S, Se, Si, Ti, Tl, V, Zn, Co and Sr), using an Optima 8300 ICP-OES inductively coupled plasma optical emission spectrophotometer (PerkinElmer) using the Waters Method. In all ICP-OES analytical runs, a Sc internal-calibration standard was continuously introduced into the plasma along with each sample, and samples were analyzed in triplicate. Quality assurance/quality control (QA/QC) samples included deionized water blanks (Barnstead Nanopure system, Thermo Fisher Scientific) that contained trace-metal-grade HNO_3_ (Thermo Fisher Scientific), and certified continuing calibration verification (CCV) standards. The QA samples were analyzed immediately after instrument calibration, after every 15 samples, and at the end of each set of samples. Additionally, NIST certified standard reference material 1643f was analyzed for trace elements before and at the end of each set of samples. All samples were reanalyzed in any analytical run in which acceptable QA/QC results were not obtained. Those unacceptable results could include deviations of the internal Sc standard greater than 20% from the known concentration, deviations of the CCV samples greater than 10% from the known concentrations, or relative standard deviations (RSDs) of triplicate analyses of a sample greater than 10%. Ion chromatography analyses were conducted on a Dionex Aquion Ion Chromatography System (ThermoScientific, Waltham, MA), consisting of a dual isocratic pump, an anion guard column (AG22 4mm, no. 064139), an anion separator column (AS22 4mm, no. 064141), coupled with an ion self-regenerating suppressor (AERS 500 Carbonate 4 mm 085029) and a DS6 Heated conductivity cell detector. Concentrations of analytes are reported in mg/L (ppm).

### Microscopy

From groundwater samples, 1 mL was removed and fixed with paraformaldehyde to a final concentration of 2% and stained with SYBR Gold. Stained cells were captured on a 0.02 µm Anodisc filter by vacuum filtration, mounted to a glass slide in the dark, and fixed with a coverslip. Cells were enumerated using a Nikon Eclipse Ti-E Widefield fluorescence microscope. A 3 x 5 grid was imaged using a 40X air objective with 300 ms exposure and 50% LED power, and a FITC filter set. Cells were manually counted on acquired images using ImageJ and converted to cell concentrations based on the volume of fluid filtered and the area of the filter. Negative controls of Anodisc filter and Anodisc Filter + MilliQ water were included. No cells were detected in these controls.

### Dissolved organic carbon (DOC) analysis of site fluids

^1^H-NMR bulk metabolite characterization was performed on aqueous organic carbon. Samples (180 μl) were combined with 2,2-dimethyl-2-silapentane-5-sulfonate-d6 (DSS-d6, Chenomx) in D_2_O (20 μl, 5 mM) and thoroughly mixed before transfer to 3 mm NMR tubes. NMR spectra were acquired on a Bruker Avance III spectrometer operating at a field strength of 17.6 T (1H ν0 of 750.24 MHz) and equipped with a 5 mm Bruker TCI/CP HCN (inverse) cryoprobe with Z-gradient and at a regulated temperature of 298 K. The one-dimensional (1D) ^1^H spectra were acquired using a Nuclear Overhauser Effect Spectroscopy (NOESY) pulse sequence (noesypr1d). The 90° ^1^H pulse was calibrated before the measurement of each sample with a spectral width of 12 ppm and 2048 transients. The NOESY mixing time was 100 ms and the acquisition time was 4 s, followed by a relaxation delay of 1.5 s during which presaturation of the water signal was applied. Time domain free induction decays (72,114 total points) were zero filled to 131,072 total points before Fourier transform, followed by exponential multiplication (0.3 Hz line-broadening) and semi-automatic multipoint smooth segments baseline correction. Chemical shifts were referenced to the 1H methyl or 13C signal in DSS-d6 at 0 ppm. The 1D ^1^H-NMR spectra of all samples were processed, assigned and analyzed using Chenomx NMR suite 9.0 (Chenomx) with quantification of spectral intensities of compounds in the Chenomx library relative to the internal standard. Candidate metabolites present in each of the complex mixtures were determined by matching chemical shift, J-coupling and intensity information of the experimental signals against signals of the standard metabolites in the Chenomx, HMDB and custom in-house databases. Signal to noise ratios (S/N) were measured using MestReNova 14.1, with the limits of quantification and detection equal to an S/N of 10 and 3, respectively.

### Dissolved inorganic carbon and δ^13^C_DIC_ measurements

Dissolved inorganic carbon concentration and carbon isotopic composition was measured in groundwater samples. Water samples and a suite of carbonate standards in 12 mL Exetainer Sample Vials (Labco) were flushed with He for 10 minutes. 1 mL of boiled water was added to standard vials and acidified with 0.2 mL of pre-boiled phosphoric acid. For mildly and moderately alkaline fluids (BA1B, BA4A), 0.5 mL sample was acidified with 0.2 mL of boiled phosphoric acid. For hyperalkaline waters (BA3A samples) exhibiting low DIC, 13 mL of site fluid was acidified with 2.6 mL of pre-boiled phosphoric acid. These were centrifuged and set on a shaker table for three days to equilibrate.

DIC and δ^13^C_DIC_ was measured on a Thermo Delta V continuous-flow isotope ratio mass spectrometer attached to a gas bench device. Data was processed using in-house scripts using R statistical software with the IsoVerse package (https://www.isoverse.org/). Stable isotope ratios are reported in delta notation relative to the Vienna Pee Dee Belemnite (VPDB) standard for carbon, where δ = ([(RSample/RStandard) − 1]), R is the ratio of the heavier mass isotope to the lighter mass isotope, as per mil (‰). δ^13^C values in this run were corrected for a small offset then scale using a 4 point scale correction. δ^13^C errors estimated using the residual standard error of the linear scale correction model.

### Gas measurements

CH_4(g)_ measurements were conducted using an SRI 8610C Gas Chromatograph coupled to a flame ionization detector (FID) equipped with a 5W amplifier. The FID gain was set to medium. Argon was used as a carrier gas at 35 psi. H_2_ at 20 psi and air at 5 psi were used for FID gases. The headspaces of the SIP incubations were anaerobically sampled using a gas-tight syringe and injected at volumes between 0.25 and 1 ml onto a 2m 2mmID ShinCarbonST 80/100 column held at 200^°^C. Methane headspace concentrations were calculated by applying a standard curve of peak area versus known methane concentrations. Total methane production was calculated as the sum of headspace (gaseous) methane and dissolved (aqueous) methane, where aqueous methane was determined using a temperature-adjusted Henry’s law. CO_2(g)_ and CO_(g)_ were also measured but not detected in any samples.

CH_4_^13^C isotopic composition was determined using a Picarro G2201-i cavity ringdown isotope analyzer. Incubation headspace was injected into a gas tight syringe filled with a known volume of ultra-pure N_2_ at atmospheric pressure (∼0.8 atm, Boulder CO). This syringe, fitted with a gas-tight septum, was attached in-line to the Picarro system, which allowed the contained gas to be pulled into the instrument. The ^13^C composition of methane was determined by averaging measured δ^13^C_VPDB_ values in high range mode (HR_Delta_iCH4_Raw) across a stable peak i.e., where ^12^CH_4_ and ^13^CH_4_ concentrations stabilized after sample injection. δ^13^C values, reported in permil, were converted to isotopic ratio (R) using the following relationship: ^*13*^*R*_*sample*_ *= ((δ*^*13*^*C*_*sample*_ *× 1000) + 1) ×* ^*13*^*R*_*VPDB*_, subsequently converted to fractional abundance (F) as ^*13*^*F =* ^*13*^*R*_*sample*_ */(1 +* ^*13*^*R*_*sample*_). Picarro CRDS data was analyzed with the IsoCRDS R package (https://github.com/KopfLab/isoCRDS). Contributions of the ^13^C label to produced methane are calculated as:

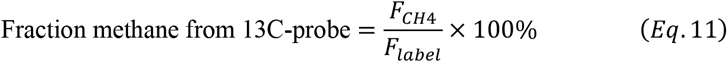

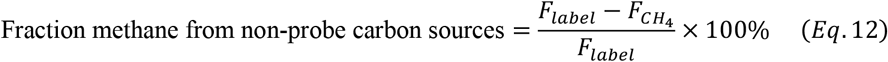

Where F represents the ^13^C isotopic fractional abundance of the measured CH_4_ and the isotopic label (F_label_). Validation of CRDS measurements at extremely high *δ*^*13*^*CH*_*4*_ was validated using an in-house calibration, described in **Supplementary Text**.

### DNA Sequencing

DNA was extracted from filters with negative controls using the DNeasy PowerSoil DNA Isolation Kit (Qiagen), according to the manufacturer’s instructions. Extracted DNA samples were amplified in duplicate using Platinum II Hot-Start PCR Master Mix (ThermoFisher Scientific) on SimpliAmp thermocycler (ThermoFisher). The 16S rRNA gene primers 515F and 806R (V4 amplification) with Illumina sequencing adapters and unique 12-bp barcodes were used as described by the Earth Microbiome Project (67). The PCR program was 94 °C for 2 min followed by 35 cycles of 94 °C (15 s), 60 °C (15 s), 68 °C (1 min), and a final extension at 72 °C for 10 min. Amplification was verified via gel electrophoresis. Amplicons were cleaned and normalized with the SequalPrep Normalization Plate (Thermo Fisher Scientific) following the manufacturer’s instructions and then pooled together. Pooled libraries were quantified using both the Qubit ds DNA High Sensitivity Assay (ThermoFisher) and the KAPA Library Quantification Kit for Illumina platform (Roche). Sequencing was performed on an Illumina MiSeq using a v2 500 cycle PE reagent kit with paired-end reads at the Center for Microbial Exploration, University of Colorado Boulder. We note that amplification bias and the difficulty of lysing archaeal cell walls may result in a decreased quantification of archaea present in this study.

To prepare samples for analysis with the DADA2 bioinformatic pipeline (68), reads were demultiplexed with adapters and primers were removed using standard settings for cutadapt. We used standard filtering parameters with slight modifications for 2 x 150 bp chemistry where forward reads were not trimmed, and reverse reads were trimmed to 140 base pairs. A maximum error rate (maxEE) of 2 was allowed. Taxonomy was assigned with the SILVA (v138) reference database (69). To control for possible contamination during DNA extraction and sample processing, we used the decontam package to filter for contaminant taxa (70). A total of 84 ASVs, prevalent in negative controls but low-abundance or absent in samples, were removed.

### Statistical Analyses

Data processing, statistical analyses, and visualization, were performed in R version 4.3.2 (71) using the *tidyverse* package suite (72). Gini coefficients were calculated and plotted with the *ineq* and *gglorenz* packages (73, 74) (**Fig. S1**). Analysis scripts, raw data, and processed data are made available in the Data Availability section.

## Data Availability

All data generated during the course of this study is available on the Open Science Framework data repository (https://osf.io/7e6ua/). All analysis scripts used to analyze and visualize data are available on GitHub (https://github.com/tacaro/Oman-2023).

## Acknowledgments

We thank Eric Ellison and Sabrina Kainz for assistance with field work preparation and sample handling. We are grateful to Sebastian Kopf, Eric Boyd, Alta Howells, Harp Batther, Eric Ellison, John Spear, Patrick Thieringer, Phyllis Lam, Michael Stocker, and Yueyue Si for valuable discussions during the preparation of this work. We thank Adam Younkin for assistance developing the CRDS-SIP calibration and Boswell Wing for access to the Picarro G2201-i instrument. We acknowledge Juerg Matter of University of Southampton for assistance with site access. 16S sequencing was supported by the University of Colorado Boulder Center for Microbial Exploration managed by Jessica Henley. Raman microspectroscopy was carried out at the CU Boulder Raman Microspectroscopy Laboratory (RRID:SCR_019305) managed by Jess Hankins and Eric Ellison. We thank and credit Dan Utter for original development of the nanoSIMS data processing script. Isotope ratio mass spectrometry was carried out at the CU Boulder Earth Systems Stable Isotope Laboratory (RRID:SCR_019300). Fluid IC and ICP-AES measurements were carried out at the Colorado School of Mines managed by Amy Ashford Boczon. Cell counts were performed using a Nikon Ti-E Widefield at the BioFrontiers Institute Advanced Light Microscopy Core (RRID: SCR_018302) managed by Joe Dragavon. The Nikon Ti-E Widefield is supported by NIH grant R01CA107098S1. DOC fluid analysis measurements were carried out at PNNL in the Environmental Molecular Sciences Laboratory, a national scientific user facility sponsored by the U.S. Department of Energy Office of Biological and Environmental Research and located on the campus of PNNL in Richland, Washington. PNNL is a multiprogram national laboratory operated by Battelle for the DOE under Contract DE-AC05-76RLO 1830. This work was supported by a NASA Exobiology Program grant to Alexis Templeton (80NSSC21K0489). Field work was supported by a Lewis and Clark Field Fund for Exploration and Field Research in Astrobiology grant to Tristan Caro. Access to the Oman Multi-borehole Observatory is thanks to the efforts of the Oman Drilling Project (https://www.omandrilling.ac.uk), NASA Astrobiology Institute Rock-Powered-Life Node, and the Oman Ministry of Regional Municipalities and Water Resources.

