## Supplemental Information for "Dissolved inorganic carbon supports robust anabolism and methanogenesis in actively serpentinizing rocks"

**This PDF file includes:**

Supplementary Text
Figures S1 to S12
Tables S1 to S8
SI References

**Other supporting materials for this manuscript include the following:**

Dataset S1

### Supplementary Text

#### Analysis of Intracommunity Heterogeneity

Single-cell estimates of microbial growth enable analyses of microbial activity heterogeneity between and within populations (1, 2). We plot the degree of intracommunity heterogeneity with Lorenz curves and represent it numerically with the Gini coefficient, a measurement of population inequality that has been adapted for descriptions of microbial communities (2). A higher Gini coefficient (closer to 1) indicates that smaller share of the microbial community is carrying out a greater share of the anabolic activity. Conversely, a lower Gini coefficient (closer to 0) indicates more even distribution of anabolic activity across a community. We observe that anabolic heterogeneity varies between each of the boreholes and depths (**Fig. S1**). Microorganisms in moderately alkaline fluids in borehole BA4A exhibited the most even anabolic activity (Gini: 0.47 – 0.58), followed by BA1B (Gini: 0.57 – 0.71). We observe the highest Gini coefficient values are observed in hyperalkaline fluids (BA3A, Gini: 0.66 – 0.93), indicating that a decreasing fraction of the microbial community is responsible for an increasing fraction of anabolic activity. We note a mild negative correlation between community-averaged microbial carbon assimilation rate and intracommunity heterogeneity measured by Gini coefficient ($F_{1, 25}$ = 15.09, $R^2$ = 0.38, p <0.001, **Fig. S1**).

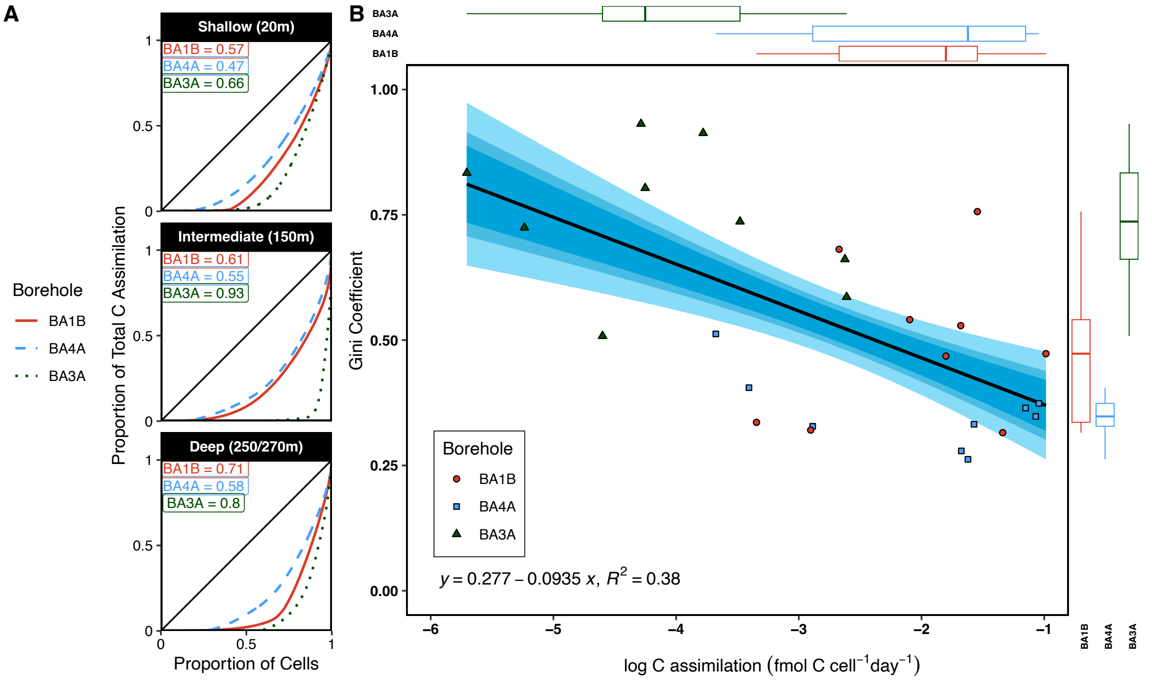


**Fig. S1. Relationship between C assimilation and intracommunity heterogeneity.**

#### Taxonomic composition of serpentinite-hosted microbial communities

The geochemically diverse groundwaters presented in this study host phylogenetically diverse microbial communities. The dominant microbial community member across all the groundwaters examined is methanogenic archaea of Genus *Methanobacterium,* as discussed in the main text. Aside from *Methanobacterium,* we observe notable differences in the non-archaeal portion of the microbial community composition between the boreholes. Mildly alkaline fluids (BA1B) contain high abundances of taxa involved in nitrogen cycling including *Azospira*, a genus of N-fixing bacteria, and *Rhodocyclaceae,* a family that includes *Azospira* as well as denitrifying bacteria including *Sulfuritalea* and *Thauera* (3). Moderately alkaline fluids in borehole BA4A contained high abundances of sulfur cycling taxa including sulfur oxidizing *Sulfuritalea*, and sulfate reducing bacteria *SRB2, Desulfonatronum, Desulfomicrobium,* and *Thermodesulfovibrionia.* This focus on sulfur cycling is likely related to higher concentrations of dissolved sulfate in groundwaters hosted in BA4A (Supplementary Data). Interestingly, we identify *Magnetospirillum*, a genus of magnetotactic bacteria, in borehole BA4A. To our knowledge, this detection, if annotated correctly, would be one of the first descriptions of this taxon in the terrestrial subsurface, possibly related to magnetite and iron cycling already observed in the system. Further targeted genomic work is required to appropriately validate this finding.

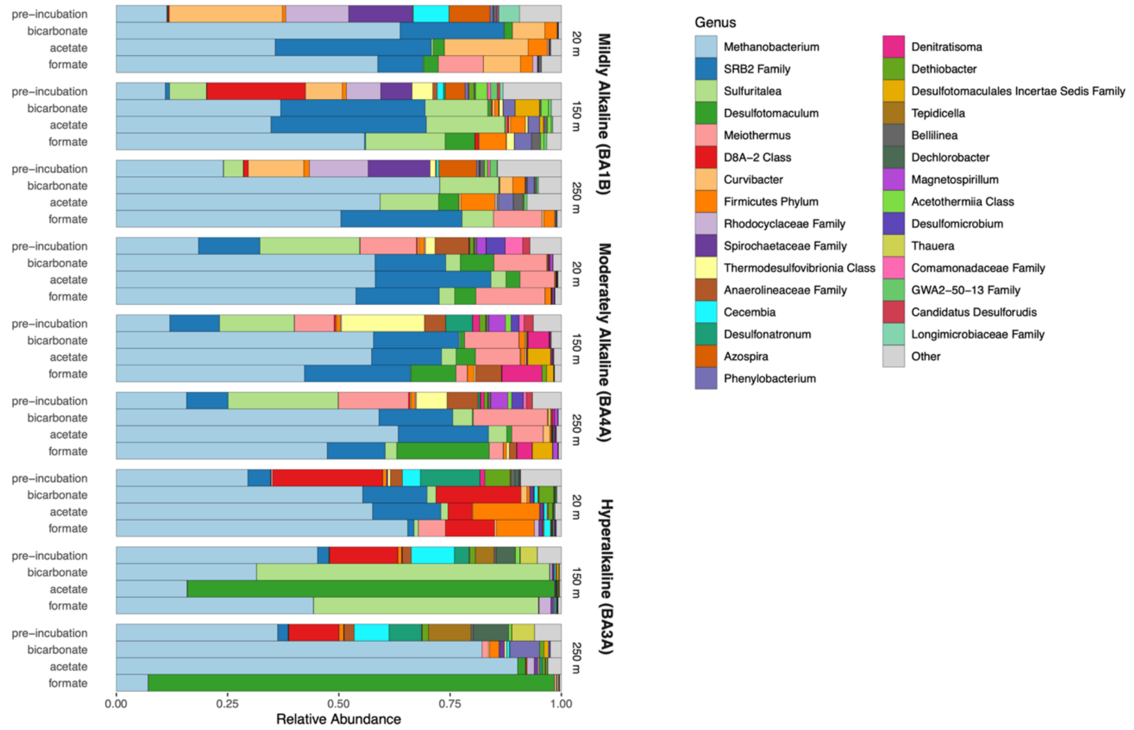


**Fig. S2. 16S rRNA gene amplicon sequencing results from borehole fluids.**
Horizontal bars indicate fractional relative abundance of 16S rRNA amplicon reads.

#### Validating CRDS-SIP measurements at high carbon isotope enrichment

Prior to measuring enriched $^{13}CH_4$ from samples on the Picarro G2201-I instrument, it was necessary to develop an in-house calibration for the instrument in order to ensure that its precision and sensitivity could span a vast potential range of $\delta^{13}CH_4$. To accomplish this, we injected methane standards of varying methane carbon isotopic composition into the instrument. Because no highly enriched of $\delta^{13}CH_4$ standards were commercially available to us, we prepared our standards in-house by careful mixing of $CH_4$ from a highly enriched source (99 at. %, Sigma Aldrich, CX1850, 490229-1L) with $CH_4$ of natural abundance (~1 at. %) isotopic composition (Airgas UHP35, Lot 49-402853298-1). Gas mixtures were prepared in gas tight syringes (SGE, Hamilton) of appropriate volumes. First, a 100 mL gas tight syringe, plugged with a GC septum (Restek) was flushed 3X with nitrogen, filled to a final volume of 95 mL, and briefly vented to equilibrate pressure with atmosphere. Natural isotopic abundance and highly enriched (99 at. %) $CH_4$ were injected at concentrations appropriate to reach the desired isotopic composition. The gases were allowed to mix in the chamber for 3 minutes at room temperature. 1 mL of the gas mixture was extracted and injected into a nitrogen-flushed 500 mL gas tight syringe (similarly sealed with a GC septum) and allowed to mix in the chamber for 3 minutes. This gas tight syringe was connected in-line to the instrument, which drew gas from the chamber for approximately 5 minutes. Results of the calibration are displayed in **Fig. S3.**

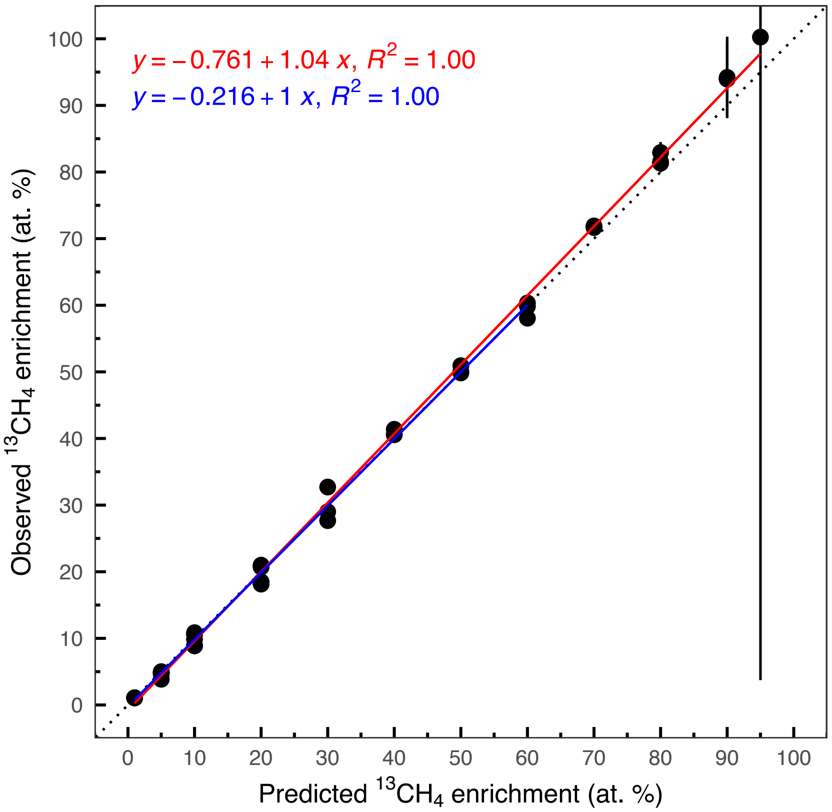


**Fig. S3.** Cavity ringdown spectroscopy (CRDS) calibration for highly enriched $^{13}CH_4$. The horizontal axis represents predicted $^{13}CH_4$ enrichment based on volumetric dilution of natural abundance $CH_4$ (~1 at. % $^{13}CH_4$) and highly $^{13}C$-enriched $CH_4$ (99 at. % $^{13}CH_4$). A linear regression from 0 – 60 at. % is plotted in blue; a linear regression across the full data range is plotted in red. The dotted line indicates a 1:1 relationship. Each point represents an individual $^{13}CH_4$ injection. Error bars indicate standard deviation in CRDS measurement ($^{13}C$ at. %) across the integrated $CH_4$ peak. Extremely high measurement error observed at 95 at. % $^{13}CH_4$ is likely due to spectral interference, but the source of this interference cannot be inferred without raw CRDS spectra (not available with this instrument). Variation between the injections is likely due to the human error associated with manual volumetric gas mixing.

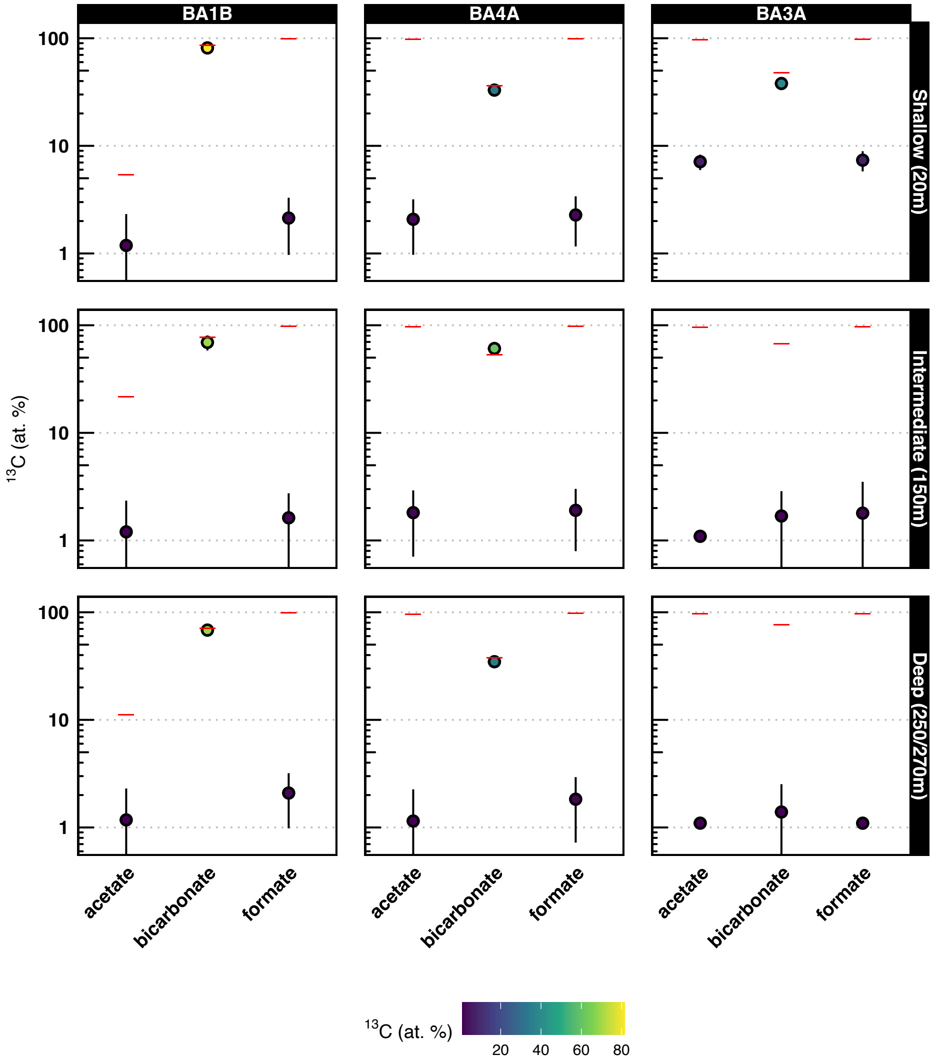


**Fig. S4. $^{13}$C enrichment of $CH_4$ measured in SIP incubation vial headspace.** The vertical axis represents $^{13}$C in fractional abundance *($^{13}F = {}^{13}C / ({}^{12}C + {}^{13}C)$)* plotted in units of atom % on a log scale. The color scale indicates $^{13}$F on a linear scale, for ease of viewing. Error bars represent standard deviation of the cavity ringdown spectroscopy data across the measured $CH_4$ peak. Horizontal red bars indicate the effective $^{13}$C tracer strength of the SIP incubation (i.e., maximum possible isotopic enrichment). Note that deep BA4A and BA3A samples were acquired at 270 m; deep BA1B samples were acquired at 250 m.

### Supplementary Figures S5 – S12

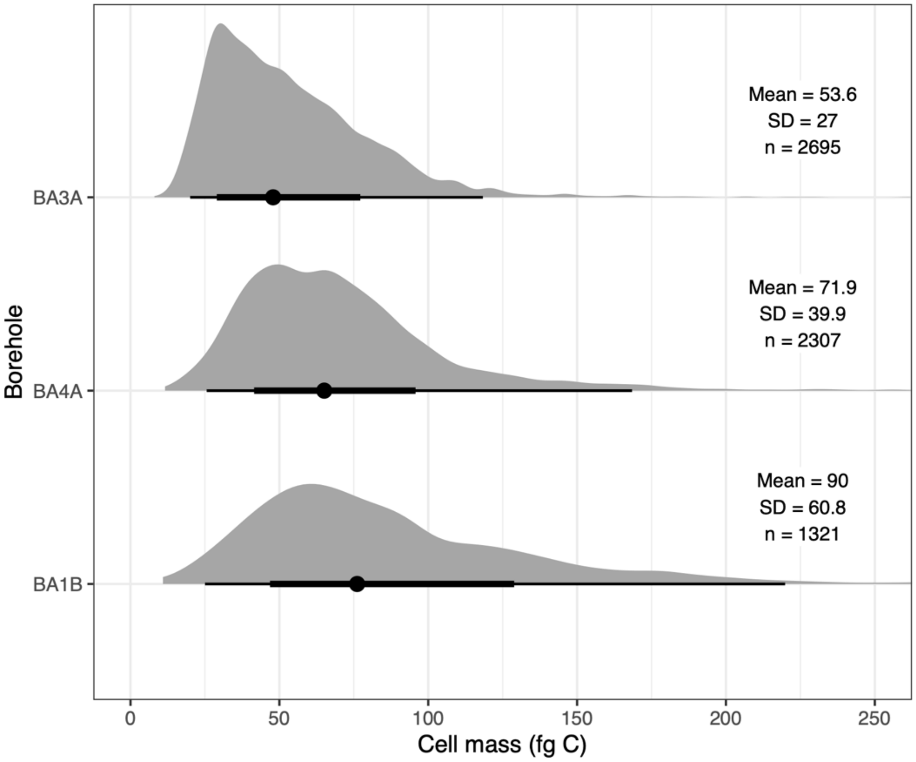


**Fig. S5.** Single-cell biomass (in fg C) estimated from nanoSIMS images across all boreholes. Data from different $^{13}$C-SIP conditions are grouped by borehole. Points mark the mean, thick bars represent 66% CI, thin bars represent 95% CI. Above the point-intervals, kernel density estimates are plotted. Deep BA4A and BA3A samples were acquired at 270 m; deep BA1B samples were acquired at 250 m.

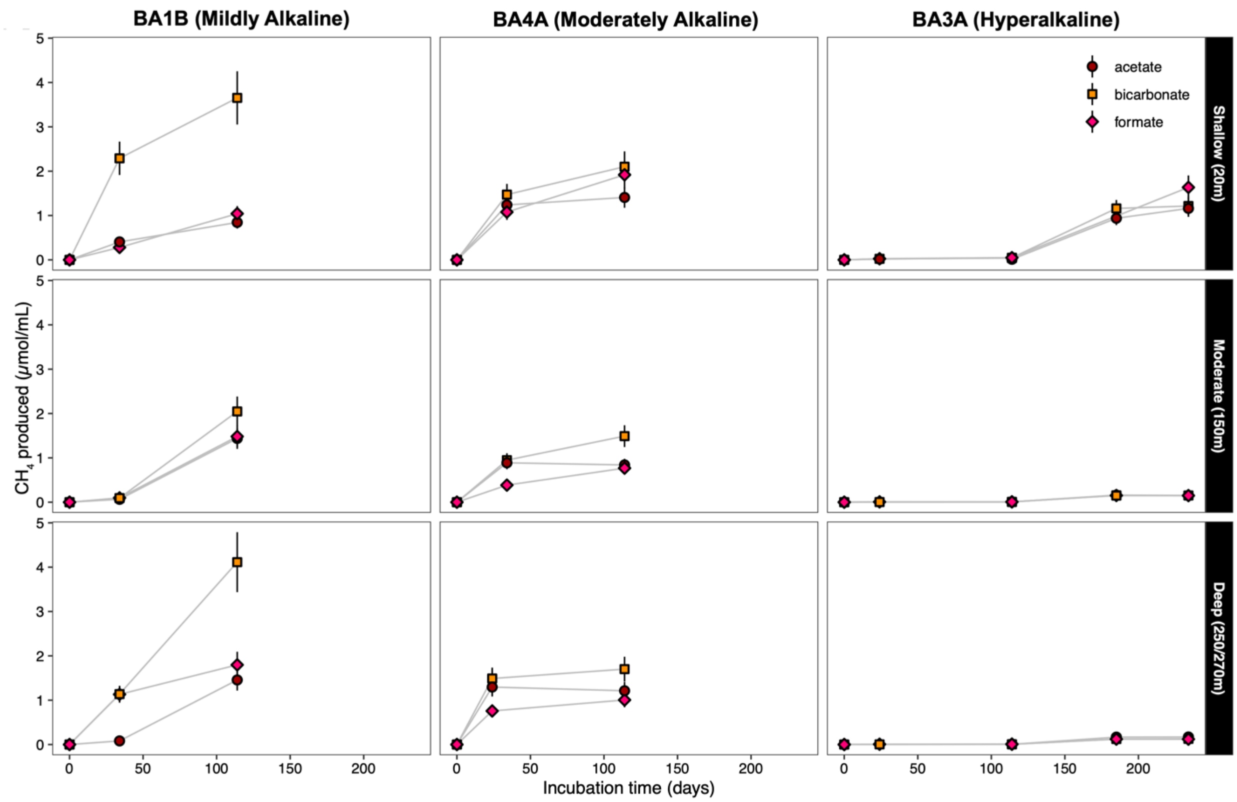


**Fig. S6.** Methane production throughout the incubation experiments as measured by GC-FID. Error bars represent the error of the standard curve calibration.

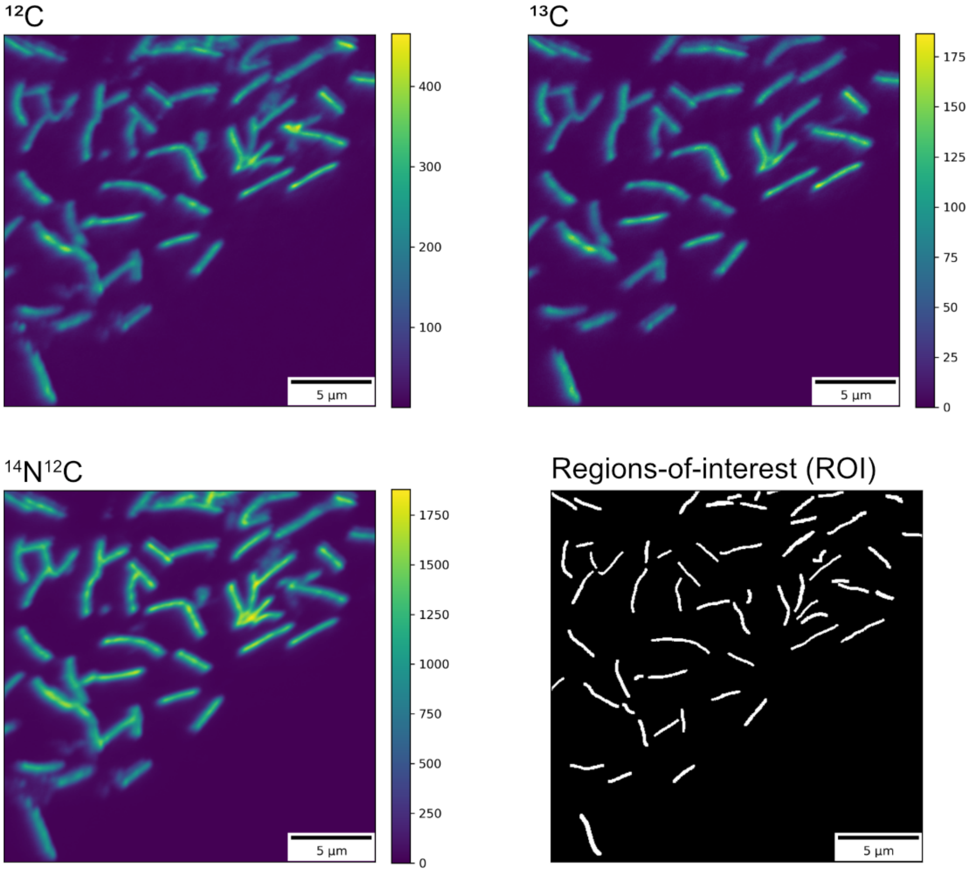


**Fig. S7. Example nanoSIMS secondary ion images.** Regions of interest (ROIs) were manually defined based on examining $^{12}C^-$, $^{13}C^-$, and $^{14}N^{12}C^-$ channels. Color bars indicate raw ion counts.

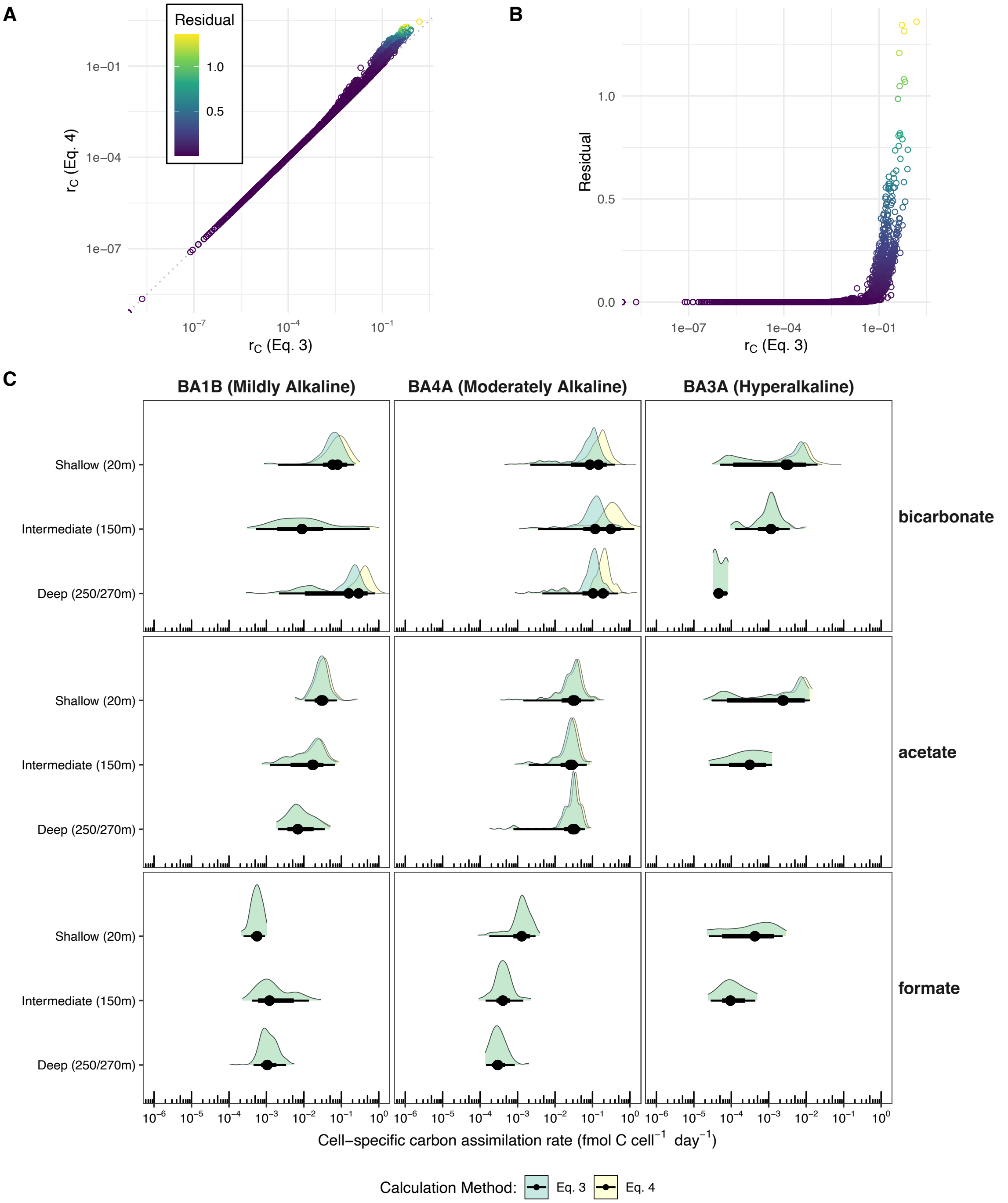


**Fig. S8. Comparison of cell-specific carbon assimilation rate methods**. As outlined in *Main Text: Materials and Methods*, we explored two different methods (Main Text: Eq. 3, Eq. 4) for estimating cell-specific carbon assimilation (4). Differences between these methods are plotted (A) directly against each other, (B) displaying residual error, and (C) in such a way as to reproduce Fig. 1. In (C), points indicate median, thick and thin bars represent 95% and 66% CI, respectively.

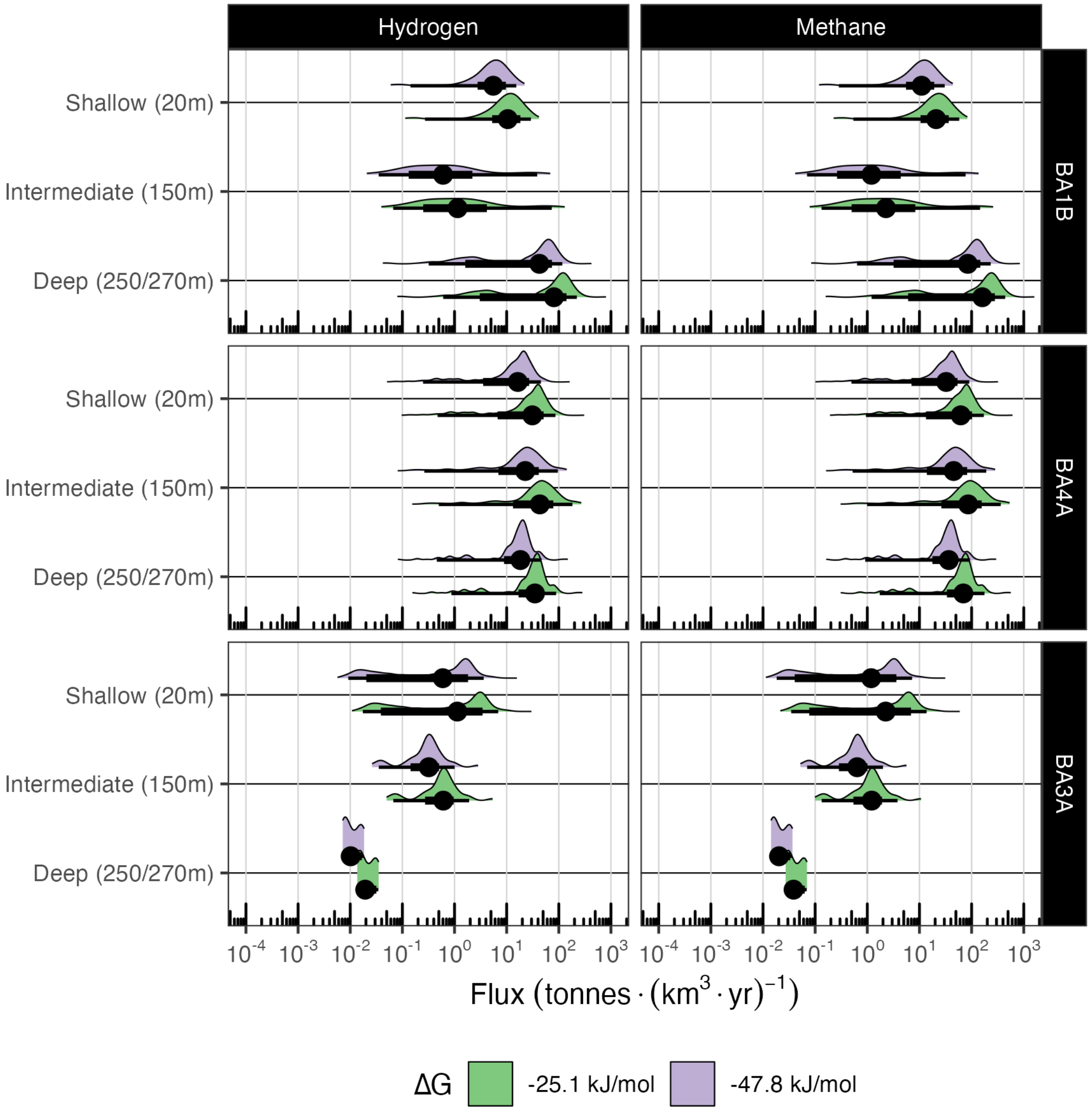


**Fig. S9. Comparison of ΔG values for calculating landscape-scale $H_2$ and $CH_4$ fluxes.** As outlined in *Main Text: Materials and Methods*, we calculated landscape-scale fluxes with two different values for Gibbs free energy (ΔG) for hydrogenotrophic methanogenesis: -25.1 kJ/mol, adapted from the range estimated by Canovas et al. (5), and -47.8 kJ/mol, which represents the standard (ΔG˚) Gibbs free energy of reaction at far from equilibrium conditions. Note that a more favorable ΔG results in lower catabolic flux, as less $H_2$ is required to account for the observed energetic requirements of carbon turnover.

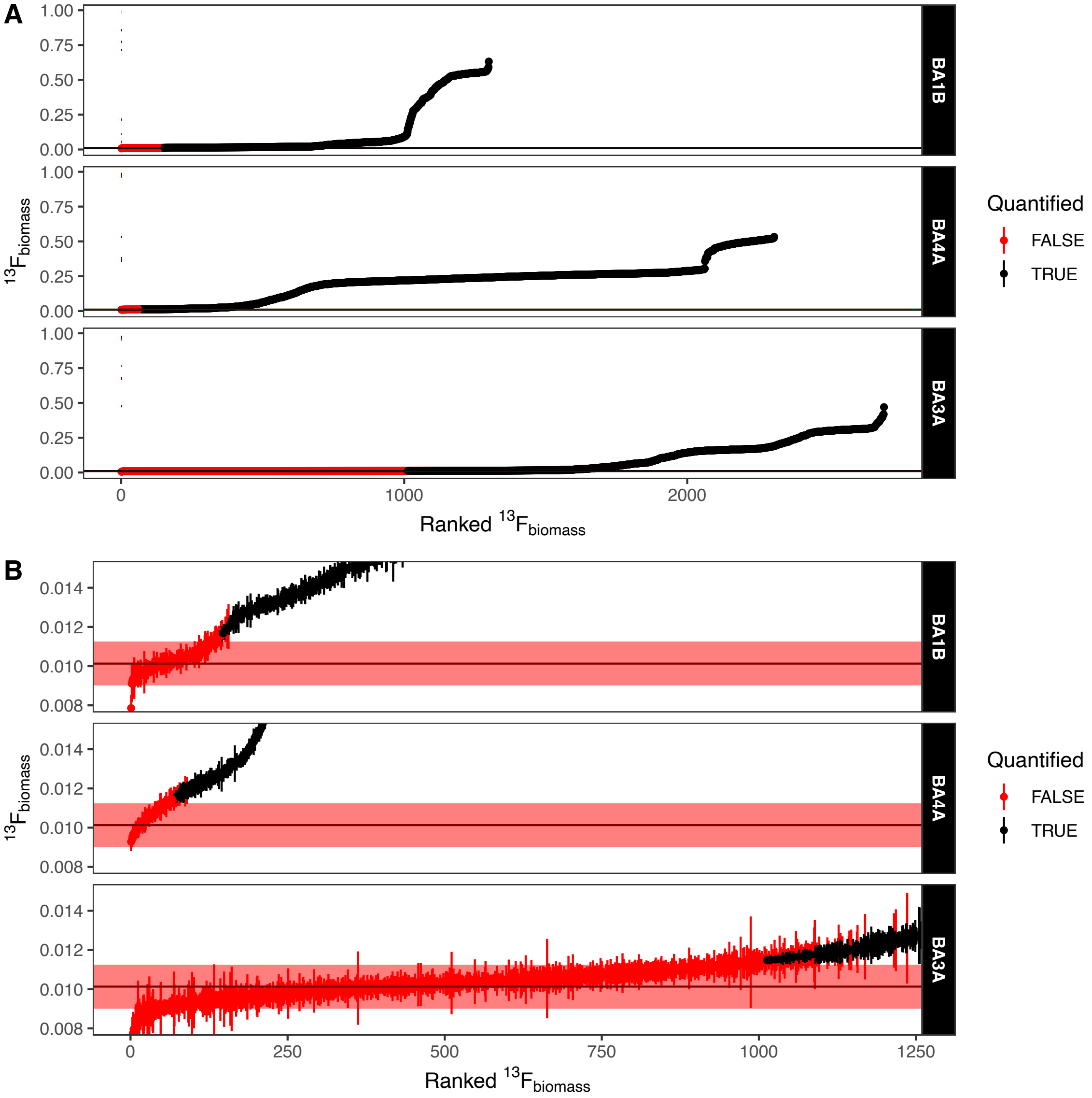


**Fig. S10. Raw $^{13}$C enrichment values across boreholes.** On both plots, the y axis indicates fractional abundance of $^{13}$C in cell biomass ($^{13}F_{biomass}$); error bar indicates the standard deviation of nanoSIMS measurement across all pixels integrated into each cell ROI. Data points are ordered in ascending order on the x-axis according to $^{13}F_{biomass}$. Panel B is a zoomed-in view of Panel A, highlighting the measurement threshold above which cells are considered "Quantified" above the negative control. The red band indicates the mean ± 3 SD of the single-cell isotopic enrichment of the negative control. Cell quantification is achieved the limit of quantification is achieved when $F_{cell} - \sigma_{cell} > F_{CTL} + (3 \times SD_{CTL})$ where $F_{cell}$ is the mean isotopic fractional abundance of a given cell region-of-interest, $\sigma_{cell}$ is the propagated error of the isotopic fractional abundance, $F_{CTL}$ is the mean isotopic fractional abundance of cells in the negative control, and $SD_{CTL}$ is the standard deviation of isotopic fractional abundance in the negative control.

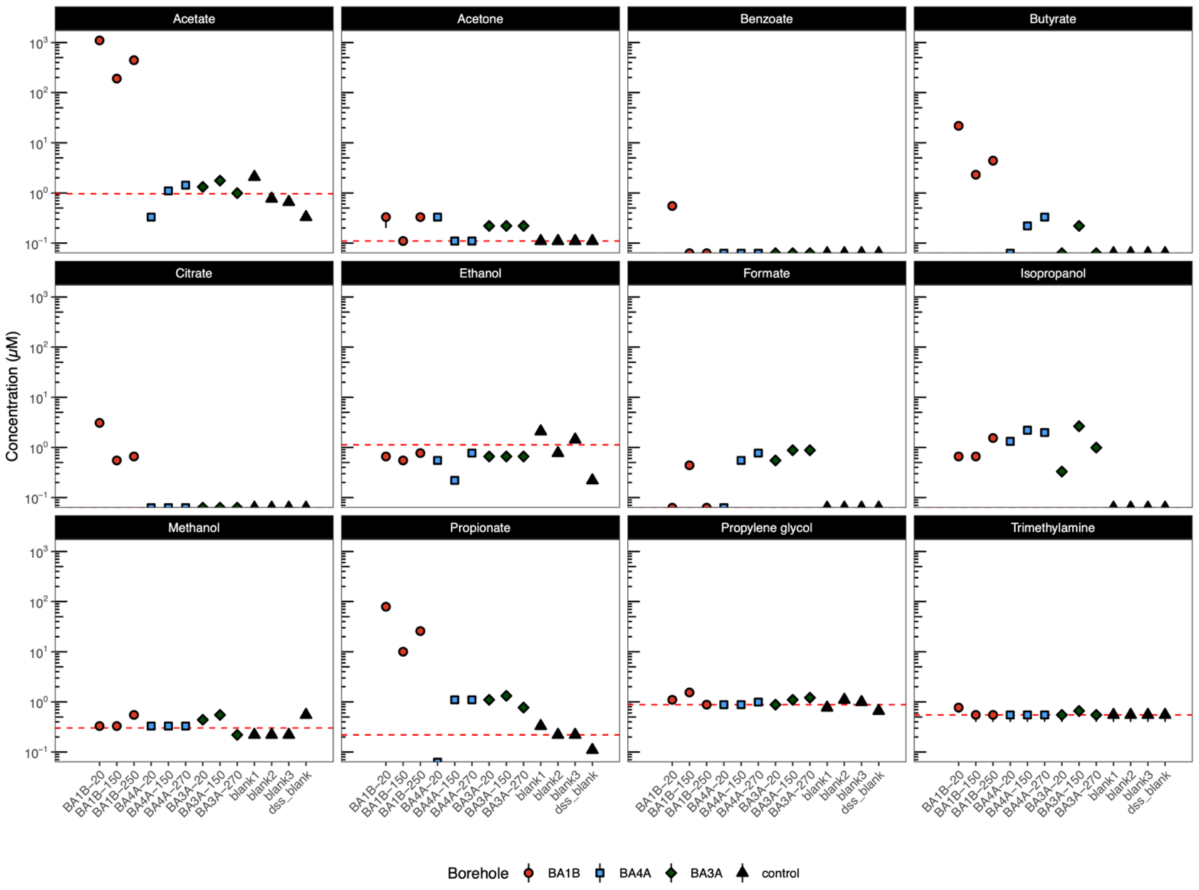


**Fig. S11. Small molecular weight organic compounds measured via liquid-state NMR.** Liquid-state NMR measurements were conducted on groundwater samples prior to incubation with $^{13}C$ probes. Concentration in µM is plotted on a logarithmic scale. The mean of the negative controls is plotted as a horizontal red line. Three processing blanks (ultrapure water), noted as blanks 1-3, and the internal standard blank "dss_blank" are plotted. Points marked on the x-axis indicate "not detected."

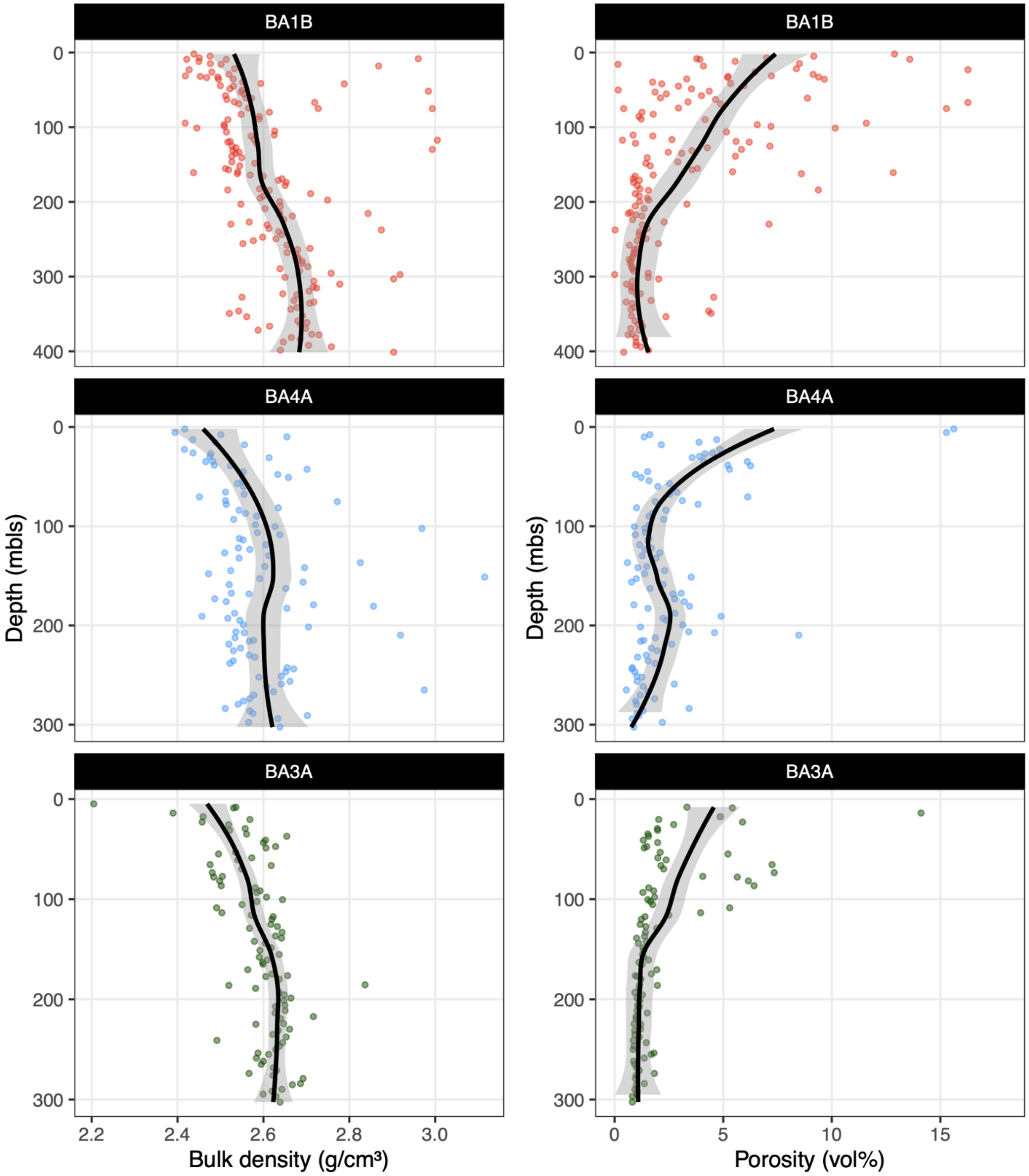


**Fig. S12.** Collated rock porosity and bulk density data reported in the *Proceedings of the Oman Drilling Project, Supplementary Information* (6) for rock cores recovered from site BA1B, BA4A, and BA3A. Line of best fit (LOESS) is shown with 95% CI. Note differing y-axes, as borehole BA1B was drilled 100 m deeper than BA4A and BA3A.

### Supplementary Tables S1 – S8

| Isotopic labeling conditions<br>Concentrations and isotopic compositions of stable isotope probes | | | | | | | | | |
|---|---|---|---|---|---|---|---|---|---|
| | Acetate | | | DIC | | | Formate | | |
| Depth (mbs) | Pre-incubation (µM) | After $^{13}C$ probe addition (µM) | Effective isotopic label (at. %) | Pre-Incubation (µM) | With $^{13}C$ Tracer (µM) | Effective isotopic label (at. %) | Pre-incubation (µM) | After $^{13}C$ probe addition (µM) | Effective isotopic label (at. %) |
| **BA1B** | | | | | | | | | |
| 20 | 1,098.02 | 1,137.04 | 5.4% | 949.38 | 6,939.88 | 85.7% | 0.00 | 50.00 | 99.0% |
| 150 | 190.63 | 238.72 | 21.6% | 1,733.84 | 7,716.50 | 77.2% | 0.44 | 50.44 | 98.2% |
| 250 | 443.74 | 489.30 | 11.1% | 2,385.68 | 8,361.83 | 71.3% | 0.00 | 50.00 | 99.0% |
| **BA4A** | | | | | | | | | |
| 20 | 0.33 | 50.33 | 98.4% | 1,802.53 | 2,784.51 | 36.3% | 0.00 | 50.00 | 99.0% |
| 150 | 1.10 | 51.09 | 96.9% | 891.50 | 1,882.58 | 53.1% | 0.55 | 50.54 | 97.9% |
| 270 | 1.43 | 51.42 | 96.3% | 1,685.99 | 2,669.13 | 37.8% | 0.77 | 50.76 | 97.5% |
| **BA3A** | | | | | | | | | |
| 20 | 1.32 | 51.31 | 96.5% | 111.53 | 210.41 | 47.6% | 0.55 | 50.54 | 97.9% |
| 150 | 1.76 | 51.74 | 95.7% | 48.20 | 147.72 | 67.4% | 0.88 | 50.87 | 97.3% |
| 270 | 0.99 | 50.98 | 97.1% | 29.87 | 129.58 | 76.7% | 0.88 | 50.87 | 97.3% |

**Table S1. Endogenous carbon sources and effective $^{13}C$ label strengths for each SIP incubation.** Cell fill is colored by relative magnitude, per column, for readability.

| Median Cell-Specific Carbon Assimilation Rates | | | | |
|---|---|---|---|---|
| | | Carbon Source | | |
| | Depth (mbls) | bicarbonate[1] | acetate[1] | formate[1] |
| BA1B (Mildly Alkaline) | 20 | $7.97 \times 10^{-2}$ | $3.20 \times 10^{-2}$ | $5.62 \times 10^{-4}$ |
| BA1B (Mildly Alkaline) | 150 | $8.97 \times 10^{-3}$ | $1.78 \times 10^{-2}$ | $1.21 \times 10^{-3}$ |
| BA1B (Mildly Alkaline) | 250 | $2.89 \times 10^{-1}$ | $6.93 \times 10^{-3}$ | $1.05 \times 10^{-3}$ |
| BA4A (Moderately Alkaline) | 20 | $1.45 \times 10^{-1}$ | $3.36 \times 10^{-2}$ | $1.29 \times 10^{-3}$ |
| BA4A (Moderately Alkaline) | 150 | $3.10 \times 10^{-1}$ | $2.89 \times 10^{-2}$ | $4.05 \times 10^{-4}$ |
| BA4A (Moderately Alkaline) | 270 | $1.91 \times 10^{-1}$ | $3.32 \times 10^{-2}$ | $2.96 \times 10^{-4}$ |
| BA3A (Hyperalkaline) | 20 | $3.30 \times 10^{-3}$ | $2.49 \times 10^{-3}$ | $4.31 \times 10^{-4}$ |
| BA3A (Hyperalkaline) | 150 | $1.16 \times 10^{-3}$ | $3.13 \times 10^{-4}$ | $9.48 \times 10^{-5}$ |
| BA3A (Hyperalkaline) | 270 | $4.61 \times 10^{-5}$ | NA | NA |

[1] All values reported in cell-specific fmol C per day.

**Table S2: Tabulated median cell-specific carbon assimilation rates.** Rates reported here are the median values (reported in cell-specific fmol C per day) from each sample site. Fill color of individual cells represents relative magnitude within each column.

| Fraction of Active Cells | | | |
|---|---|---|---|
| | BA1B | BA4A | BA3A |
| **Shallow (20m)** | | | |
| acetate | 65.55% | 100.00% | 88.91% |
| bicarbonate | 97.86% | 99.56% | 73.17% |
| formate | 80.23% | 94.74% | 53.26% |
| **Medium (150m)** | | | |
| acetate | 89.71% | 100.00% | 13.06% |
| bicarbonate | 98.65% | 100.00% | 13.02% |
| formate | 74.42% | 78.57% | 37.04% |
| **Deep (250 / 270m)** | | | |
| acetate | 76.85% | 100.00% | 0.00% |
| bicarbonate | 99.54% | 100.00% | 24.14% |
| formate | 97.44% | 50.00% | 0.00% |

**Table S3.** Fraction of cells determined to be actively assimilating the indicated carbon source. Cells are color-coded by borehole.

| Depth (mbls) | Median MSP (W/g C) |
|---|---|
| BA1B | |
| 20 | $2.54 \times 10^{-3}$ |
| 150 | $6.05 \times 10^{-4}$ |
| 250 | $5.88 \times 10^{-4}$ |
| BA4A | |
| 20 | $9.36 \times 10^{-3}$ |
| 150 | $2.69 \times 10^{-3}$ |
| 270 | $2.53 \times 10^{-3}$ |
| BA3A | |
| 20 | $3.09 \times 10^{-4}$ |
| 150 | $5.65 \times 10^{-5}$ |
| 270 | $8.52 \times 10^{-6}$ |

**Table S4.** Median MSP in each borehole, across all carbon sources tested. Fill color of individual cells represents relative magnitude.

| Depth (mbls) | Hydrogen Consumed, (tonnes $yr^{-1}$ $km^{-1}$) | Methane Produced, (tonnes $yr^{-1}$ $km^{-1}$) |
|---|---|---|
| BA1B | | |
| 20 | $1.05 \times 10^{1}$ | $2.09 \times 10^{1}$ |
| 150 | 1.15 | 2.28 |
| 250 | $8.06 \times 10^{1}$ | $1.60 \times 10^{2}$ |
| BA4A | | |
| 20 | $3.11 \times 10^{1}$ | $6.18 \times 10^{1}$ |
| 150 | $4.32 \times 10^{1}$ | $8.59 \times 10^{1}$ |
| 270 | $3.49 \times 10^{1}$ | $6.94 \times 10^{1}$ |
| BA3A | | |
| 20 | 1.13 | 2.25 |
| 150 | $6.10 \times 10^{-1}$ | 1.21 |
| 270 | $1.94 \times 10^{-2}$ | $3.85 \times 10^{-2}$ |

**Table S5**. Estimates of $H_2$ oxidation and $CH_4$ production at the reservoir scale based on single-cell MSP estimates. Cells are color-coded by the relative magnitude of the enclosed value, per column.

| Borehole | Depth (mbls) | Sampling Device | Temperature (˚C) | Dissolved $H_2$ (µM) | Dissolved $H_2$ (µM) [Corrected][1] |
|---|---|---|---|---|---|
| WAB188 | 50 | open-borehole pumping | 35.3 | $6.62 \times 10^{-1}$ | $2.29 \times 10^{1}$ |
| BA1A | 250 | open-borehole pumping | 36.6 | $1.83 \times 10^{-1}$ | 6.31 |
| BA1D | 50 | open-borehole pumping | 34.6 | $4.13 \times 10^{-2}$ | 1.43 |
| BA1B | 50 | open-borehole pumping | 33.7 | $5.06 \times 10^{-3}$ | $1.75 \times 10^{-1}$ |
| BA4A | 50 | open-borehole pumping | 36.1 | $9.82 \times 10^{1}$ | $3.40 \times 10^{3}$ |
| BA4A | 80 | open-borehole pumping | 36.9 | $1.40 \times 10^{2}$ | $4.84 \times 10^{3}$ |
| BA3A | 55 | open-borehole pumping | 36.2 | 4.82 | $1.67 \times 10^{2}$ |
| BA3A | 75 | open-borehole pumping | 36.8 | $4.18 \times 10^{1}$ | $1.45 \times 10^{3}$ |
| BA3A | 50 | gas-tight tool | 35.4 | $1.67 \times 10^{2}$ | $1.67 \times 10^{2}$ |
| BA3A | 100 | gas-tight tool | 36.4 | $5.87 \times 10^{2}$ | $5.87 \times 10^{2}$ |
| BA3A | 275 | gas-tight tool | 40.8 | $1.47 \times 10^{3}$ | $1.47 \times 10^{3}$ |

[1] Corrected by estimated dilution factor (34.5X) from open-borehole pumping relative to gas-tight tool.

**Table S6.** Dissolved $H_2$ concentrations previously measured at the Samail Ophiolite during the 2020 field session (February – March 2020). Note differences in $H_2$ concentrations dependent on sampling device employed: open-borehole pumping via submersible pump (typically discharging at rates of multiple L/min) versus a discrete interval, hermetically-sealed gas-tight sample retrieval tool. Comparing $H_2$ concentrations from BA3A 55 m and 50 m, sampled with open-borehole pumping and the gas-tight tool, respectively, provides a rough dilution factor estimate for the effect of submersible pumping on measurable $H_2$ concentrations. Boreholes BA1A and BA1D are nearby BA1B, described in this study, whereas WAB188 is distant from the "BA" boreholes (7, 8).

| date | Depth, mbls | Sampling Device | pH | Conductivity, µS $cm^{-1}$ | Temperature, °C | Depth to water table, mbls |
|---|---|---|---|---|---|---|
| BA1B | | | | | | |
| 2023-01-25 | 20 | Point-Source Bailer | 7.83 | 498 | 23.1 | 18 |
| 2023-01-23 | 80 | Open-Borehole Submersible Pump | 7.87 | 450 | 35.1 | 18 |
| 2023-01-25 | 150 | Point-Source Bailer | 8.94 | 368 | 27.1 | 18 |
| 2023-01-25 | 250 | Point-Source Bailer | 8.08 | 491 | 24.8 | 18 |
| BA4A | | | | | | |
| 2023-01-26 | 20 | Point-Source Bailer | 9.22 | 491 | 25.3 | 13.6 |
| 2023-01-24 | 80 | Open-Borehole Submersible Pump | 9.45 | 533 | NA | 13.6 |
| 2023-01-26 | 150 | Point-Source Bailer | 9.91 | 604 | 27.8 | 13.6 |
| 2023-01-26 | 270 | Point-Source Bailer | 9.54 | 507 | 27.2 | 13.6 |
| BA3A | | | | | | |
| 2023-01-24 | 20 | Point-Source Bailer | 10.32 | 1439 | 28.4 | 8 |
| 2023-01-25 | 80 | Open-Borehole Submersible Pump | 11.41 | 3500 | 36.9 | 8 |
| 2023-01-24 | 150 | Point-Source Bailer | 11.44 | 2960 | 27.2 | 8 |
| 2023-01-24 | 270 | Point-Source Bailer | 11.72 | 3330 | 25.7 | 8 |
| NSHQ14 | | | | | | |
| 2023-01-26 | 80 | Open-Borehole Submersible Pump | 11.34 | 2850 | 36.1 | NA |

**Table S7. Groundwater characteristics measured on-site.** Note that temperature measured at the point-source bailer is lower than actual subsurface temperatures because of the time it took samples to be collected allowed water to cool. Subsurface fluid temperature is more faithfully represented by groundwaters collected via Open-Borehole Submersible Pump, around 35-36˚C, **(Table S6)**.

| Borehole | Depth (mbls) | $^{13}C$ tracer | Incubation Time (days) |
|---|---|---|---|
| BA1B | All | All | 40 |
| BA4A | All | All | 40 |
| BA3A | 20 | Acetate, bicarbonate, formate | 238 |
| BA3A | 150 | Acetate, bicarbonate | 246 |
| BA3A | 270 | Acetate | 246 |
| BA3A | 270 | Bicarbonate, formate | 254 |

**Table S8**. **SIP Incubation durations.** Differences in incubation duration result from time needed for adequate metabolic activity to be observed (*Main Text: Materials and Methods*) as well as time required for sample preparation and quality control via Raman microspectroscopy.

**Dataset S1 (separate file).**

Dataset S1 contains supplementary data (Excel workbook format, .xlsx) referenced in this manuscript.
